# Timing the SARS-CoV-2 Index Case in Hubei Province

**DOI:** 10.1101/2020.11.20.392126

**Authors:** Jonathan Pekar, Michael Worobey, Niema Moshiri, Konrad Scheffler, Joel O. Wertheim

**Author notes:** Corresponding authors. (MW); (JOW).

## Abstract

Understanding when SARS-CoV-2 emerged is critical to evaluating our current approach to monitoring novel zoonotic pathogens and understanding the failure of early containment and mitigation efforts for COVID-19. We employed a coalescent framework to combine retrospective molecular clock inference with forward epidemiological simulations to determine how long SARS-CoV-2 could have circulated prior to the time of the most recent common ancestor. Our results define the period between mid-October and mid-November 2019 as the plausible interval when the first case of SARS-CoV-2 emerged in Hubei province. By characterizing the likely dynamics of the virus before it was discovered, we show that over two-thirds of SARS-CoV-2-like zoonotic events would be self-limited, dying out without igniting a pandemic. Our findings highlight the shortcomings of zoonosis surveillance approaches for detecting highly contagious pathogens with moderate mortality rates.

## Introduction

In late-December of 2019, the first cases of COVID-19, the disease caused by SARS-CoV-2, were described in the city of Wuhan in Hubei province, China (*1, 2*). The virus quickly spread within China (*3*). The cordon sanitaire that was put in place in Wuhan on 23 January 2020 and mitigation efforts across China eventually brought about an end to sustained local transmission. In March and April 2020, restrictions across China were relaxed (*4*). By then, however, COVID-19 was a pandemic (*5*).

A concerted effort has been made to determine when the virus first began transmitting among humans by retrospectively diagnosing the earliest cases of COVID-19. Both epidemiological and phylogenetic methods have been used to suggest an emergence of the pandemic in Hubei province at some point in late-2019 (*2, 6, 7*). The first described cluster of COVID-19 was associated with the Huanan Seafood Wholesale Market in late-December 2019, and the earliest sequenced SARS-CoV-2 genomes came from this cluster (*8, 9*). However, this market cluster could not have marked the beginning of the pandemic, as COVID-19 cases from early December lacked connections to the market (*6*). The earliest such case in the scientific literature is from an individual retrospectively diagnosed on 01 December 2019 (*7*). Notably, however, newspaper reports document retrospective COVID-19 diagnoses recorded by the Chinese government going back to 17 November 2019 in Hubei province (*10*). In fact, these reports detail daily retrospective COVID-19 diagnoses through the end of November, suggesting that SARS-CoV-2 was actively circulating for at least a month before it was discovered.

Molecular clock phylogenetic analyses have inferred the time of most recent common ancestor (tMRCA) of all sequenced SARS-CoV-2 genomes to be in late November or early December 2019, with uncertainty estimates typically ranging back to October 2019 (*6, 11, 12*). Crucially, though, this tMRCA is not necessarily equivalent to the date of zoonosis or index case infection (*13*), because coalescent processes can prune basal viral lineages before they have the opportunity to be sampled, potentially pushing SARS-CoV-2 tMRCA estimates forward in time from the index case by days, weeks, or months (Fig. 1). For a point of comparison, consider the zoonotic origins of the HIV-1 pandemic, whose tMRCA in the early 20^th^ century coincides with the urbanization of Kinshasa, in the Congo (*14, 15*), but whose cross-species transmission from a chimpanzee reservoir occurred in southeast Cameroon, likely predating the tMRCA of sampled HIV-1 genomes by many years (*16*). Despite this important distinction, the tMRCA has been frequently conflated with the date of the index case infection in the SARS-CoV-2 literature (*6, 17, 18*).

**Fig. 1.**
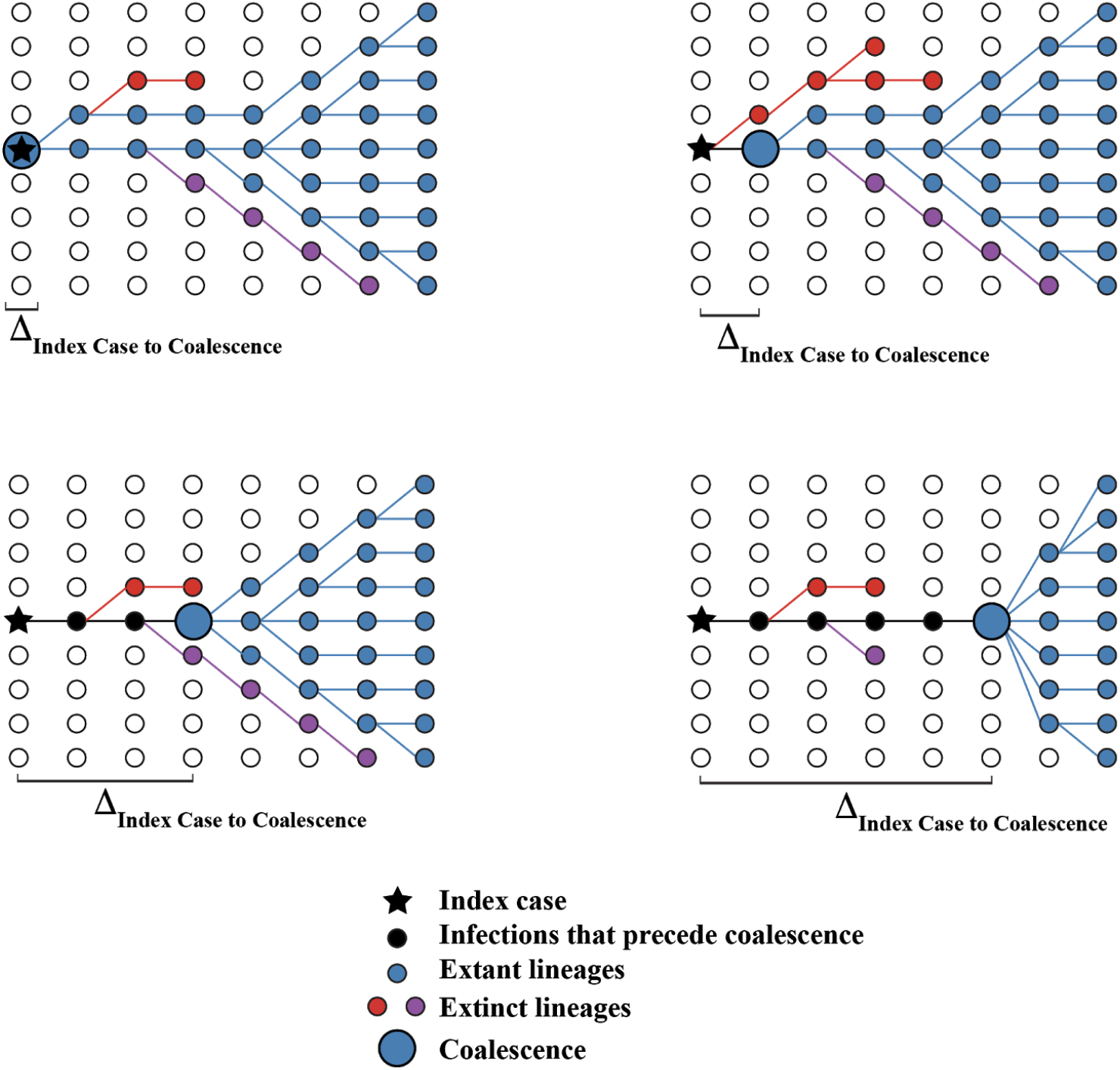
Hypothetical coalescent scenarios depicting how the time between index case infection and time of stable coalescence can vary based on stochastic extinction events of basal viral lineages. Coalescence can occur within the index case (upper left) or in cases infected later in the course of the epidemic. In extreme cases, the epidemic can persist at low levels for a long time before coalescence (lower right).

Here, we combine retrospective molecular clock analysis in a coalescent framework with a forward compartmental epidemiological model to estimate the timing of the SARS-CoV-2 index case in Hubei province. The inferred dynamics during the early days of SARS-CoV-2 highlight challenges in detecting and preventing nascent pandemics.

## Results

### Time of the most recent common ancestor of SARS-CoV-2 in China

We first explored the evolutionary dynamics of the first wave of SARS-CoV-2 infections in China. We used a Bayesian phylodynamic approach (*19*) that reconstructed the underlying coalescent processes for 583 SARS-CoV-2 complete genomes, sampled in China between when the virus was first discovered at the end of December 2019 and the last of the non-reintroduced circulating virus in April 2020. Applying a strict molecular clock, we inferred an evolutionary rate of 7.90×10^-4^ substitutions/site/year (95% highest posterior density [HPD]: 6.64×10^-4^-9.27×10^-4^). The tMRCA of these circulating strains was inferred to fall within a 34-day window with a mean of 07 December 2019 (95% HPD: 17 November–20 December) (Fig. 2). This estimate accounts for many disparate rooting orientations inferred here (see Supplementary Material) and described previously (*20*). Notably, 78.7% of the posterior density post-dates the earliest published case on 01 December, and 95.1% post-dates the earliest reported case on 17 November. Relaxing the molecular clock provides a similar tMRCA estimate (Fig. S1). The recency of this tMRCA estimate in relation to the earliest documented COVID-19 cases obliges us to consider the possibility that this tMRCA does not capture the index case and that SARS-CoV-2 was circulating in Hubei province prior to the inferred tMRCA.

**Fig. 2.**
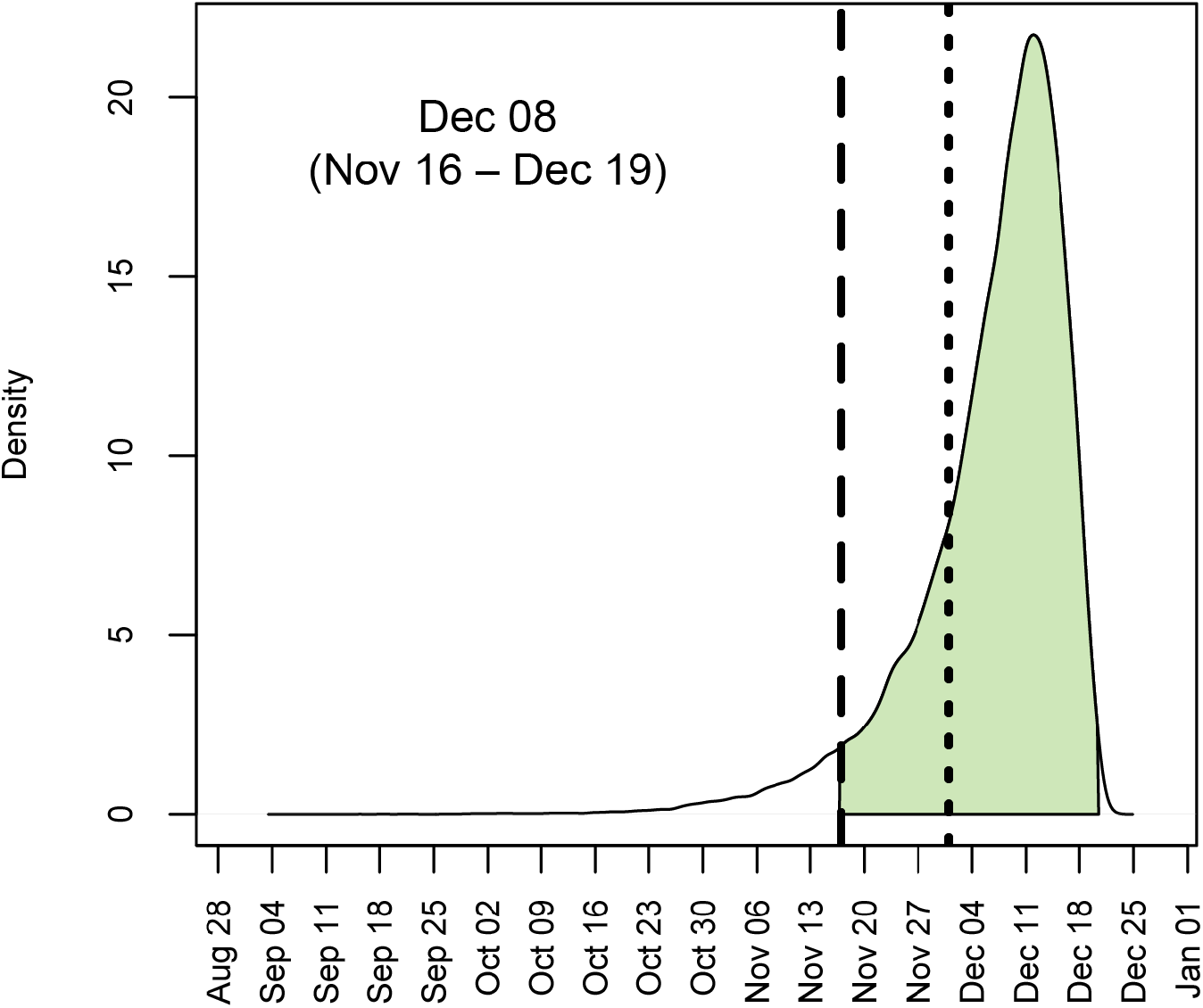
Posterior distribution for the time of the most recent common ancestor (tMRCA) of 583 sampled SARS-CoV-2 genomes circulating in China between December 2019 and April 2020. Shaded area denotes 95% HPD. Long-dashed line is 17 November 2019, and short-dashed line is 01 December 2019.

### Continued loss of basal lineages

If the tMRCA post-dates the earliest documented cases, then the earliest diverged SARS-CoV-2 lineages must have gone extinct. As these early basal lineages disappeared, the tMRCA of the remaining lineages would move forward in time (Fig. S2). Thus, we interrogated the posterior trees sampled from the phylodynamic analysis to determine if this time of coalescence had stabilized prior to the sequencing of the first SARS-CoV-2 genomes on 24 December 2019 or if this process of basal lineage loss was ongoing in late-December/early-January. Importantly, these basal lineages need not be associated with specific mutations, as the phylodynamic inference reconstructs the coalescent history, not the mutational history (*19*).

We find only weak evidence for basal lineage loss between 24 December 2019 and 13 January 2020 (Fig. S3A). The root tMRCA is within 1 day of the tMRCA of virus sampled on or after 01 January 2020 in 78.5% of posterior samples (Fig. S3B). The tMRCA of genomes sampled on or after 01 January 2020 is 3 days younger than the tMRCA of all sampled genomes. In contrast, the mean tMRCA does not change when considering genomes sampled on or after 01 January 2020 versus on or after 13 January 2020. This consistency indicates a stabilization of coalescent processes at the start of 2020 when there had been an estimated 1000 total people infected with SARS-CoV-2 in Wuhan (*22*). Nonetheless, to account for the weak signal of a delay in reaching a stable coalescence, we identified the tMRCA for all viruses sampled on or after 01 January 2020 (i.e., at the time of stable coalescence) for each tree in the posterior sample.

### Simulating the Wuhan epidemic

Phylogenetic analysis alone cannot tell us how long SARS-CoV-2 could have circulated in Hubei province before the tMRCA. To answer this question, we performed forward epidemic simulations (*21*). These simulations were initiated by a single index case using a compartmental epidemiological model across scale-free contact networks (mean number of contacts=16). This compartmental model was previously developed to describe SARS-CoV-2 transmission dynamics in Wuhan (*22*). This model, termed SAPHIRE, includes compartments for susceptible (S), exposed (E), presymptomatic (P), unascertained (A), ascertained (I), hospitalized (H), and removed (R) individuals. Our simulations used parameters from the time-period prior to COVID-19 mitigation efforts, from 01 January through 22 January 2020 (Table S1), based on Hao et al. (*22*). We analyzed 1000 epidemic simulations that resulted in ≥1000 total infected people. These simulated epidemics had a median doubling time of 4.1 days (95% range across simulations: 2.7-6.7), matching pre-mitigation incidence trends in Wuhan (Table S2).

We simulated coalescent processes across the transmission network to determine the tMRCA of the virus at the end of the simulation. This approach allowed us to determine the distribution of the expected number of days between index case infection and the stable coalescence: the tMRCA (Fig. 1). The median number of days between index case infection and this tMRCA was 8.0 days (95% range: 0.0 and 41.5 days) (Fig 3a). The median time between index case infection and the first person exiting the presymptomatic phase (i.e., ascertained or unascertained infection) was 5.7 days (95% range: 0.9 to 15.7 days).

**Fig. 3.**
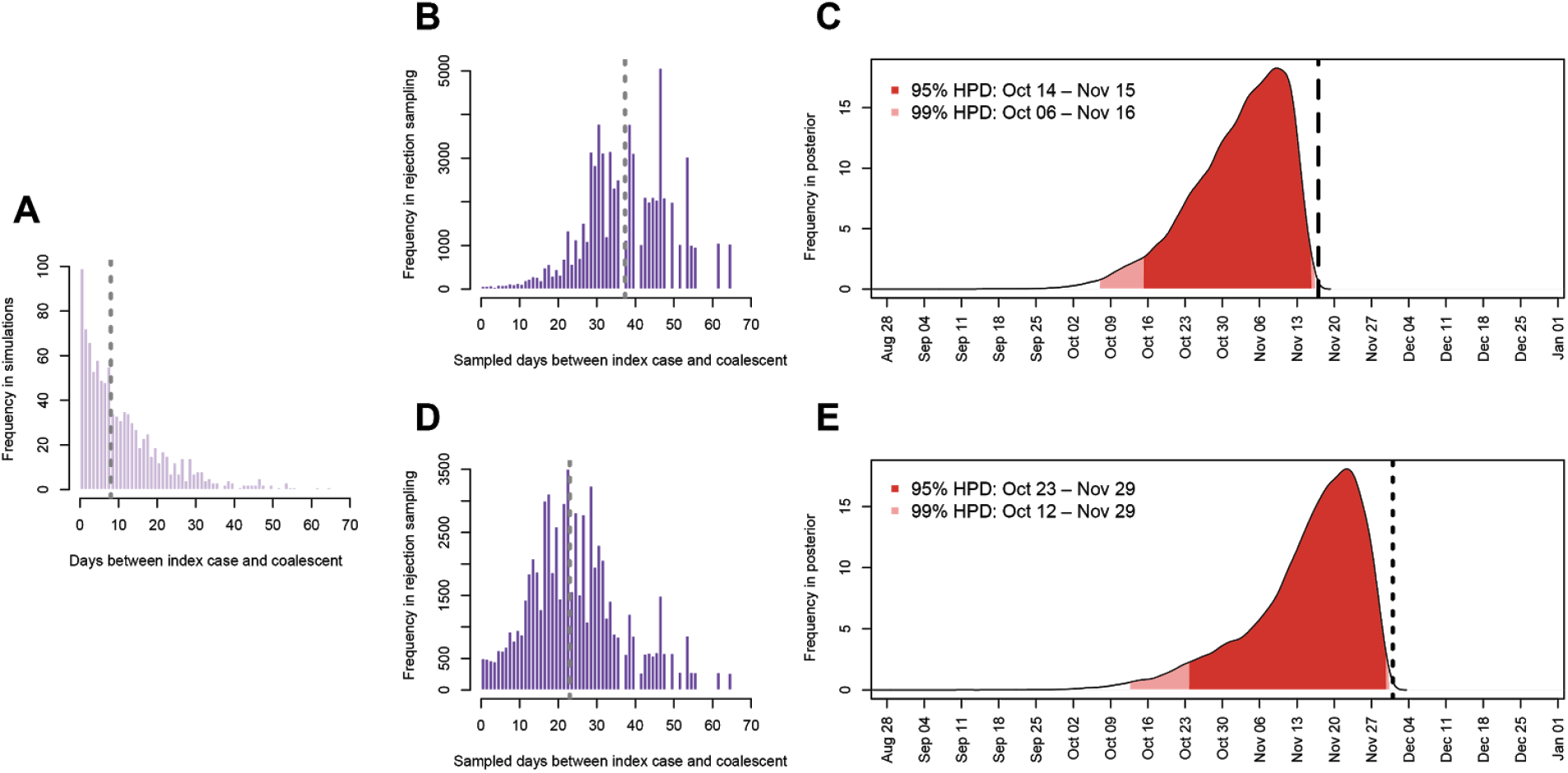
Forward simulations estimating the timing of the index case in Hubei province. (A) Days between index case infection and stable coalescence in forward compartmental epidemic simulations (n=1000). (B) Days between index case infection and stable coalescence after rejection sampling, conditioned on an ascertained case by 17 November 2019. (C) Posterior distribution for date of index case infection, conditioned on an ascertained case by 17 November 2019 denoted by long-dashed line. (D) Days between index case infection and stable coalescence after rejection sampling, conditioned on an ascertained case by 01 December 2019. (E) Posterior distribution for date of index case infection, conditioned on an ascertained case by 01 December 2019 denoted by short-dashed line. Grey dashed lines indicate median estimates.

As a robustness check, we also simulated epidemics with greater and fewer mean number of contacts in the contact network (26 and 10 contacts, respectively). We also explored the effects of faster (mean: 3.1 days; 95% range: 2.0 to 5.1 days) and slower (mean: 5.3 days; 95% range: 3.6 to 7.5 days) epidemic doubling times (Table S2). Slower transmission rates led to more days between the index case and the stable coalescence, but the connectivity of the contact network had minimal effect (Fig. S4).

### Timing of the Hubei Index Case

To estimate the date of infection for the index case in Hubei province, we combined the retrospective molecular clock analysis with the forward epidemic simulations (Fig. 4). We identified the stable tMRCA in the posterior trees as an anchor to the real-world calendar dates and then extended this date back in time according to the number of days between the index case infection and the time of stable coalescence from the compartmental epidemic simulations. However, a random sample of tMRCAs and days from index case infection to coalescence will not produce epidemiologically meaningful results, because many of these combinations do not precede the earliest dates of reported COVID-19 cases. Therefore, we implemented a rejection sampling approach to generate a posterior distribution of dates of infection for the Hubei index case, conditioning on at least one individual who has progressed past the pre-symptomatic stage in the simulated epidemic before the date of the first reported COVID-19 case (see Methods; Fig. S5).

**Fig. 4.**
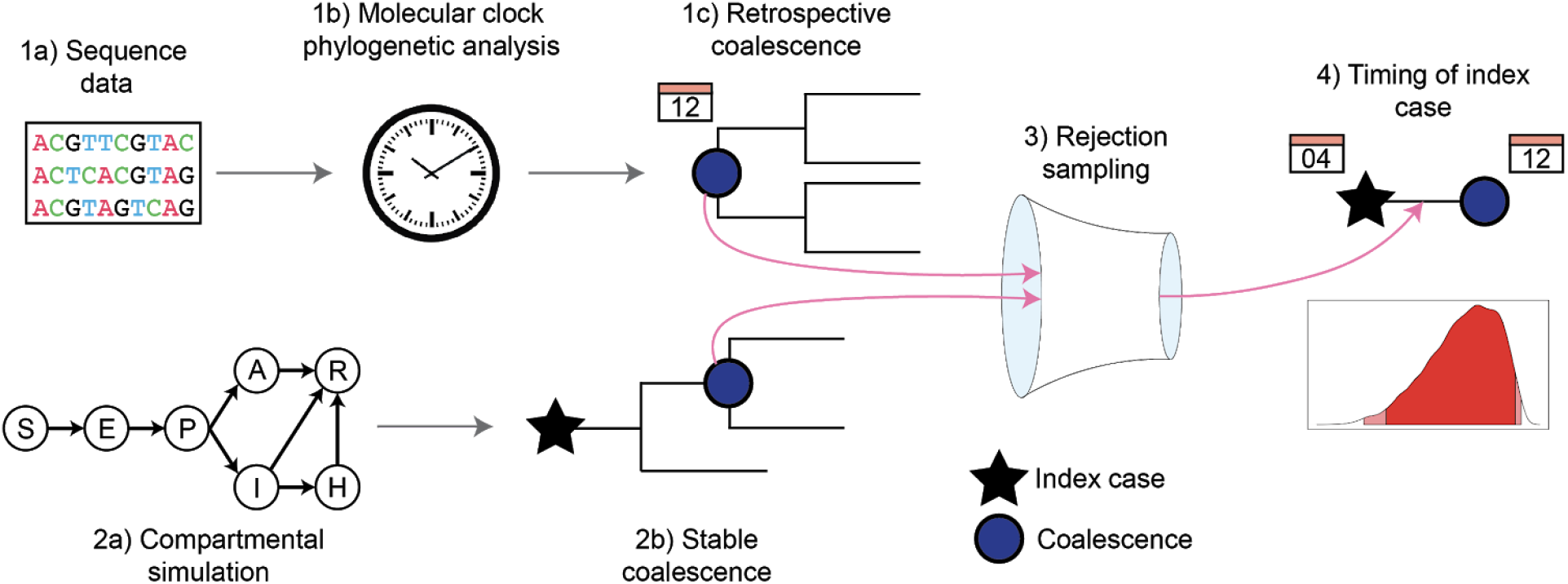
Combined simulation and phylogenetic workflows to estimate the timing of the Hubei index case. (1a) Using sequence and epidemiological data, (1b) BEAST performs a phylodynamic molecular clock analysis to (1c) determine the tMRCA. (2a) FAVITES simulates the epidemic in Hubei using a SAPHIRE compartmental model (*22*) and (2b) estimates a prior distribution for the time from index case to the stable coalescence. The results of (*1*) and (*2*) are combined via rejection sampling (3; Fig. S5) to (*4*) determine the timing of the index case and its posterior distribution.

In our primary analysis, we assume that 17 November represents the first documented case of COVID-19 (ascertained or unascertained in the SAPHIRE model). Under this assumption, the median number of days between index case infection and stable coalescence after rejection sampling is 37 days (95% HPD: 12 to 55 days) (Fig 3b). Consequently, the index case in Hubei likely contracted SARS-CoV-2 on or around 04 November 2019 (95% upper HPD on 16 October; 99% upper HPD on 07 October) (Fig 3c).

This timeframe for the Hubei index case is robust (Fig. S6). Epidemic simulations with faster or slower transmission rates and more or less densely connected contact networks produce similar date estimates. Further, using the root tMRCA from the sampled posterior trees, rather than adjusting for the shifting coalescence between 24 December 2019 and 01 January 2020, produces a similar distribution: median date 02 November (95% upper HPD 13 October; 99% upper HPD 03 October). Incorporating a relaxed molecular clock into the phylodynamic analysis shifts this distribution only slightly further back in time relative to the strict clock: median date 02 November (95% upper HPD 15 October; 99% upper HPD 02 October).

If we enforce that there be at least one ascertained case in our simulations prior to 17 November 2019, the median date of the Hubei index case is pushed back about a week to 27 October (95% upper HPD 10 October; 99% upper HPD 04 October) (Fig. S6). However, the distinction between ascertained and unascertained in the original SAPHIRE model was meant to reflect the probability of missed diagnoses in January 2020 of the Wuhan epidemic and does not account for the investigations that resulted in retrospective diagnoses in November and December 2019.

If we discount the reported evidence of retrospective COVID-19 diagnoses throughout the end of November and instead take 01 December as representing the first confirmed case of COVID-19, then the median time between index case infection and stable coalescence after rejection sampling is 23 days (95% range: 1 to 47 days) (Fig 3d). Under this scenario, the index case in Hubei would have contracted SARS-CoV-2 on or around 27 November 2019 (95% upper HPD 24 October; 99% upper HPD 13 October) (Fig 3e). Similar dates are inferred with a relaxed molecular clock and conditioning on an ascertained infection by 01 December (Fig. S6).

### Expected number of cases early in the outbreak

By anchoring our epidemic simulations to specific tMRCA estimates, we can reconstruct a plausible range for the number of SARS-CoV-2 infections prior to the discovery of the virus (Fig. 5A). The median number of individuals infected with SARS-CoV-2 in our simulations is below 1 until 04 November. On 17 November, the median number infected is 4 individuals (95% HPD: 1-13) and reaches 9 (95% HPD: 2-26) on 01 December. These values are generally robust to model specifications, molecular clock method, and date of first COVID-19 case (Table S3; Fig. S7). Further, we do not see any evidence for an increase in hospitalizations until mid-to-late December (Supplementary Material; Fig. S8).

**Fig. 5.**
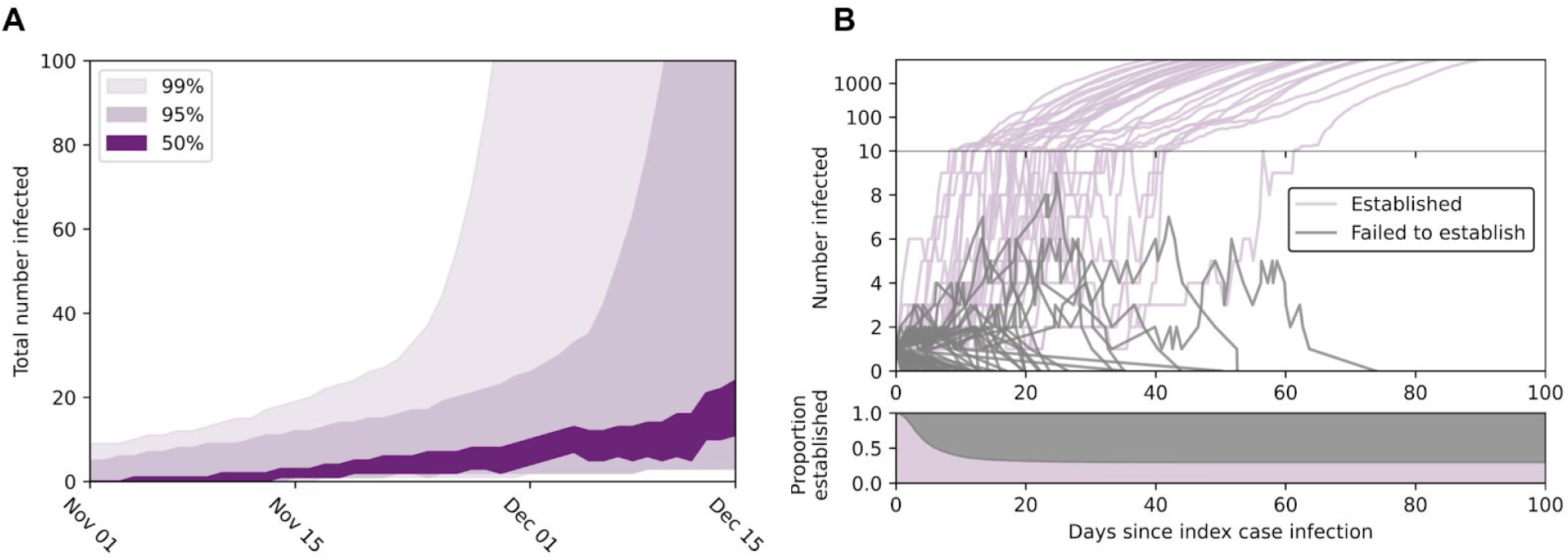
Epidemic growth in compartmental simulations. (A) Total estimated people infected in late-2019. Dark purple shading is central 50% HPD, intermediate purple shading is central 95% HPD, and light purple is central 99% HPD. (B) The number of people infected over time in a sample of epidemic simulations that established (purple; n=30) and failed to establish (grey; n=70). The y-axis transitions to log-scale once ≥10 people are infected at any given time. The lower panel shows the proportion of simulations that still have ≥1 infected individual over time (persisting epidemics in purple; extinct epidemics in grey).

### Expected number of mutations separating the index case and root

It has been speculated that if SARS-CoV-2 was circulating in humans prior to the tMRCA, this time-period could have permitted the evolution of human-specific viral adaptations, specifically in the polybasic cleavage site in the Spike protein (*13*). Based on posterior estimates for the tMRCA, substitution rate, and number of days between index case infection and the tMRCA, we estimate that approximately 2.2 mutations (95% HPD: 0.5-3.9) occurred in SARS-CoV-2 before giving rise to the observed patterns of genetic diversity (*23*). This estimate is similar if we assume a relaxed molecular clock: 2.5 mutations (95% HPD: 0.1-4.3).

### Probability of establishing a self-sustaining epidemic

Empirical observation throughout the SARS-CoV-2 pandemic has shown the outsized role of superspreading events in the propagation of SARS-CoV-2 (*24–27*), wherein the average infected person does not transmit the virus. Our results suggest the same dynamics likely influenced the initial establishment of SARS-CoV-2 in humans, as only 29.7% of simulated epidemics went on to establish self-sustaining epidemics. The remaining 70.3% of epidemics went extinct (Fig. 5B).

Epidemics that went extinct typically produced only 1 infection (95% range: 1-9) and never more than 44 infections total or 14 infections at any given time (Table S3). The median failed epidemic went extinct by day 8. The number of epidemics that went extinct was similar as the contact network became more or less densely connected, 68.3% and 69.4%, respectively. However, the percentage of extinct epidemics increased as the transmission rate decreased (80.5%) and decreased as the transmission rate increased (53.6%).

## Discussion

Our results highlight the unpredictable dynamics that characterized the earliest days of the COVID-19 pandemic. The successful establishment of SARS-CoV-2 post-zoonosis was far from certain, as more than two-thirds of simulated epidemics quickly went extinct. It is highly probable that SARS-CoV-2 was circulating in Hubei province at low levels in early-November 2019 and possibly as early as mid-October 2019, but not earlier. Nonetheless, the inferred prevalence of this virus was too low to permit its discovery and characterization for weeks or months. By the time COVID-19 was first identified, the virus had firmly established itself in Wuhan. This delay highlights the difficulty in surveillance for novel zoonotic pathogens with high transmissibility and moderate mortality rates.

The same dynamics that characterized the establishment of SARS-CoV-2 in Hubei province may have played out all over the world as the virus was repeatedly introduced, but only occasionally took hold (*28, 29*). The existence of early cases reported in December 2019 and January 2020 in France and California that did not establish sustained transmission fit this pattern (*30–32*). However, our results suggest that PCR evidence of SARS-CoV-2 in sewage outside of China before November 2019 is unlikely to be valid (*33*) and the suggestion of international spread in late-November or early-December 2019 should be viewed with skepticism (*34, 35*), given that our results suggests fewer than 20 people infected with SARS-CoV-2 at this time (Table S3; Fig. S7). On the other hand, SARS-CoV-2 may be detectable should archived waste water samples from Hubei province exist from early-to-mid November 2019. This approach may present the best chance of early detection of future, similar pandemics during the early phase of spread where we estimate very low numbers of infections. Our results also refute claims (*36*) of large numbers of patients requiring hospitalization due to COVID-19 in Hubei province prior to December 2019 (Fig. S8).

Our dating inference is insensitive to geography: even though all of the earliest documented cases of COVID-19 were found in Hubei province, we cannot discount the possibility that the index case initially acquired the virus elsewhere. However, the lack of reports of COVID-19 elsewhere in China in November and early-December suggest Hubei is the first location where human-to-human transmission chains were first established.

Finding the animal reservoir or hypothetical intermediate host for SARS-CoV-2 could help to further narrow down the date, location, and circumstances of the original SARS-CoV-2 infection in humans. However, even in the absence of that information, coalescent-based approaches permit us to look back beyond the tMRCA and towards the earliest days of the COVID-19 pandemic. Although there was a pre-tMRCA fuse to the COVID-19 pandemic, it was almost certainly very short. This brief period of time suggests that this pandemic, like potential future ones with similar characteristics, permitted only a narrow window for preemptive intervention.

## Methods

### Sequence data

We queried the GISAID database SARS-CoV-2 viral genome alignment for sequences from mainland China that were not annotated as travel associated, as of 16 July 2020. We restricted our dataset to genomes that (i) were complete ≥29,000 nt, (ii) had high coverage with ≤0.5% unique amino acid mutations, (iii) had fewer than 1% ‘N’s, (iv) were not identified as potentially problematic via NextStrain, and (v) had a day-month-year sampling date reported. The first 200 and last 299 nucleotides were removed due to poor evidence of homology. The final alignment comprised 583 taxa. List of GISAID ids are available at https://github.com/pekarj/SC2_Index_Case.

### Phylogenetic inference

The phylogenetic history of SARS-COV-2 in China was first inferred in a maximum likelihood framework in IQ-TREE 2 (*37*) using a GTR+F+I model, selected by model testing. Molecular clock analysis was conducted using a Bayesian Markov chain Monte Carlo (MCMC) approach in BEAST v1.10.4 (*19*). For the primary analysis, we employed a GTR+F+I substitution model, a strict molecular clock, a Bayesian skyline coalescent prior. To facilitate convergence, (i) a hard lower bound of 1×10^-5^ substitutions/site/year was placed on the clock rate and (ii) we initiated the MCMC using the maximum likelihood phylogeny that had been transformed into a chronogram via TempEst v1.5.3 (*38*). Four independent chains of 500 million generations were run, sampling every 25 thousand, and the first 15% were discarded as burnin. Convergence and mixing was assessed in Tracer v1.7.1 (*39*) and chains were combined in LogCombiner, such that all ESS values were >200. The resulting posterior distribution comprised over 70,000 sampled trees to facilitate fuller exploration in the rejection sampling (see below). Evidence for shifting root tMRCAs after excluding the earliest sampled SARS-CoV-2 genomes was explored using TreeStat, a part of the BEAST package. Robustness analysis was conducted using a relaxed clock with a uncorrelated lognormal distribution (ULD). We did not explore the effect of highly structured coalescent priors (e.g., constant size, exponential growth), because the skyline dynamics from China depict a complicated history with stable, exponential, and decreasing effective population sizes (Fig. S9).

### Epidemic Simulation

To explore the evolutionary dynamics at play during the beginning of the COVID-19 pandemic, we performed a series of epidemic simulations using FAVITES v1.2.6 (*21*) First, we generated static contact networks in FAVITES under a preferential-attachment model using the Barabási-Albert algorithm (*40*). We used this network algorithm, because its scale-free properties recapitulate infectious disease spread. We chose to simulate a static contact network, because our focus is on the number of people infected at the beginning of the epidemic. For the primary simulations, we selected an intermediate value of 16 contacts per day (mean degree), based on Mossong et al. (*41*), within a contact network comprising 100,000 individuals (nodes).

Across this contact network, we performed a forward simulation SAPHIRE (Susceptible-Ascertained-Presymptomatic-Hospitalized-Not Ascertained (I)-Removed-Exposed) model (*22*) to generate a viral transmission network using GEMF (*42*). We did not include the travel component of the original SAPHIRE model (i.e., individuals flying into and out of Wuhan), because our focus was on the early dynamics of the pandemic before its spread. Simulated epidemics started with a single seed infection among our 100,000 susceptible individuals. The epidemic was propagated using the parameters determined by Hao et al. (*22*) (see Table S3 for values) for 100 days. Our epidemics had a median doubling time of 4.1 days (95% range: 2.7-6.7 days), corresponding to epidemic growth in Wuhan between 01 January 2020 and 23 January 2020 (*22*).

For the primary analysis, we ran 5000 epidemic simulations. Of these simulations, 29.7% successfully established epidemics, defined as those simulations in which ≥1000 people had become infected and ≥1 person was still infectious at the end of the simulation. Failed epidemics were simulations that did not become established (i.e., 0 infectious people at the end of the simulation) or had fewer than 1000 people infected over the entire simulation; 70.3% of simulations failed to reach this epidemic threshold after 100 days. For the successfully established simulations, we constructed the coalescent history using the Virus Tree Simulator package in FAVITES. For each infected individual in these simulations, a single viral lineage was randomly sampled uniformly across the duration of infection to represent viral genotype sampling. If a simulation failed to coalesce, it was rerun until it successfully achieved coalescence. The final output generated by FAVITES is the viral time-based phylogeny (Fig. 4). 1000 successfully established simulations were then randomly selected for further analysis. All inputs for the FAVITES primary analysis can be found in Table S4 and the JSON input files are available at https://github.com/pekarj/SC2_Index_Case.

Each FAVITES simulation with the SAPHIRE model produced an output documenting when individuals transitioned from one compartment to another throughout the entire simulation. We used these to determine the amount of individuals in a given compartment (e.g., total infections, ascertained infections, unascertained infections, and hospitalized individuals) across each day in the simulation.

### Determining Stable Coalescence

Once we had the time-based phylogenies, we labeled the internal nodes using the FAVITES helper script *label_internal_nodes.py*. We then extracted the tMRCA of infected individuals every day across each simulation using TreeSwift 1.1.14 (*43*). This tMRCA was calculated for each day of the 100 days or until 10,000 individuals had been infected, whichever came first. We chose not to explore dynamics after 10,000 infections due to a slowing in exponential growth arising from the saturation of the contact network.

We defined the stable coalescence as the coalescence that does not shift forward in time by more than one day, even as new individuals become infected and previously infected individuals recover. Therefore, the stable coalescence is reached the first day that the coalescence for the currently infected individuals is within one day of the time of coalescence after the 100 day simulation or once 10,000 total individuals have been infected.

### Sensitivity Analysis—Faster rate of infection

The same methods for epidemic simulations were performed to evaluate the dynamics of a more rapidly spreading virus. We used the aforementioned parameters, except the edge-based rates of transmission were increased (Table S2). We produced 5000 simulations to generate at least 1000 established replicates with at least 1000 infected individuals each. Increasing the infectiousness coefficient to 0.55 per day produced a median epidemic doubling time of 3.1 days (95% range: 1.6-5.1).

### Sensitivity Analysis—Slower rate of infection

The same methods for epidemic simulations were performed to evaluate the dynamics of a more slowly spreading virus. We used the aforementioned parameters, except the edge-based rates of transmission were increased (Table S2). We produced 7000 simulations to generate at least 1000 established replicates with at least 1000 infected individuals each. Decreasing the infectiousness coefficient to 0.3 per day produced a median epidemic doubling time of 5.3 days (95% range: 3.6-7.5) (Table S2).

### Sensitivity Analysis—Higher average degree

The same methods for epidemic simulations were performed to evaluate the dynamics of a more densely connected network. We used the aforementioned parameters, except the average degree of the contact network was 26, and we adjusted the SAPHIRE parameters accordingly (Table S2). We produced 5000 simulations to generate at least 1000 established replicates with at least 1000 infected individuals each.

### Sensitivity Analysis—Lower average degree

The same methods for epidemic simulations were performed to evaluate the dynamics of a less densely connected network. We used the aforementioned parameters, except the average degree of the contact network was 10, and we adjusted the SAPHIRE parameters accordingly (Table S2). We produced 5000 simulations to generate at least 1000 established replicates with at least 1000 infected individuals each.

### Combining FAVITES and BEAST via Rejection Sampling

Our aim is to obtain a posterior distribution for the date *X* of the index case in Hubei province, conditioned on both the available sequencing data *D_s_* and the date of the first reported COVID-19 case *D_c_*. We do this in a Bayesian framework by marginalizing over the date *Y* of the first simulated COVID-19 case (either ascertained or unascertained, see below) and the date *Z* of the tMRCA as follows:

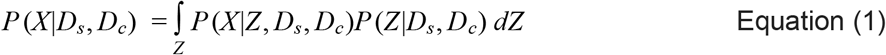

We assume that the sequencing data are informative only for the tMRCA, i.e. given *Z, X* does not depend on *D_s_*: *P*(*X|Z, D_s_, D_c_*) = *P*(*X|Z, D_c_*). We also assume that the first reported COVID-19 case data are not informative for the tMRCA: *P*(*Z|D_s_, D_c_*) = *P*(*Z|D_s_*). This gives:

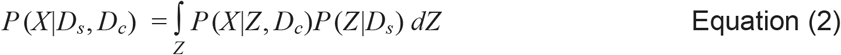

We further note that 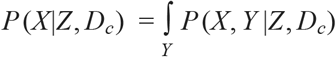, where we model *P*(*X, Y|Z, D_c_*) as proportional to *C*(*Y*)*P*(*X, Y|Z*), where *C*(*Y*) is a “consistency function” with a value of 1 when *Y* is consistent with *D_c_* and 0 otherwise. This approach allows us to sample from the posterior distribution of Equation 2. The BEAST analysis provides values of *Z* sampled from the distribution *P*(*Z|D_s_*). For each sampled value of *Z*, we sample corresponding values of *X* and *Y* from the distribution *P*(*X, Y|Z, D_c_*) using the FAVITES simulation (providing samples from the distribution *P*(*X, Y|Z*) in conjunction with a simple rejection sampling strategy: sample values from *P*(*X, Y|Z*) until a sample is obtained for which *C*(*Y*) = 1. The resulting set of sample values for *X* then follow the posterior distribution *P*(*X|D_s_, D_c_*).

We require the first simulated case to be ascertained (SAPHIRE stage: I) or unascertained (SAPHIRE stage: A) (I/A: Fig. S5) and assign *D_c_* as 17 November 2019. However, we note that this first ascertained/unascertained case can be the index case themselves, unless a secondary or tertiary case progresses faster through the course of infection. Importantly, the rate at which cases were ascertained in the SAPHIRE model is based on real-time patterns in COVID-19 diagnosis from 01 through 22 January 2020 and may not reflect the actions that led to the retrospective diagnosis of earliest cases of COVID-19. Further, coalescence can happen any time after the index case is first infected, and there is no requirement for coalescence to occur after the first ascertained and unascertained individuals.

### Sensitivity Analysis—ascertained and unascertained cases

The primary rejection sampling analysis was conditioned on an individual having an ascertained (SAPHIRE stage: I) or unascertained (SAPHIRE stage: A) infection prior to date of the first reported case of COVID-19 on 17 November 2019. However, we also performed sensitivity analyses whereby the minimum date for the earliest case must be an ascertained case (Fig. S5). We performed rejection sampling using the first ascertained or unascertained case (I/A; Fig. S5) or solely the first ascertained case (I-only; Fig. S5).

### Sensitivity Analysis—date of first COVID-19 case

We also explored the sensitivity of the rejection sampling approach to the date before which an ascertained or unascertained case must have existed. We performed rejection sampling using 17 November 2019 or 01 December 2019 as the minimum date for the earliest case.

### Sensitivity Analysis—ULD clock

We explored the sensitivity of the rejection sampling approach to different molecular clocks in the BEAST inference. We performed separate rejection sampling analyses using tMRCAs inferred under a strict clock and a relaxed ULD clock.

### Sensitivity Analysis—shifting the tMRCA

We explored the sensitivity of the rejection sampling approach to the stability of the timing of coalescence in the BEAST analysis (Fig. S3A). The primary analysis used the inferred tMRCA of all viruses sampled on or after 01 January 2020, at which point the tMRCA had stabilized. Sensitivity analysis was conducted using the tMRCA of all sampled viruses (i.e., the root tMRCA). This tMRCA is within 1 calendar day in 78.5% of sampled trees (Fig. S3B) and represented by the same node in 74.6% of sampled trees (Fig. S3C).

### Mutation analysis

To approximate the number of mutations that separated the index case virus from that represented by the most recent common ancestor of all 583 analyzed SARS-CoV-2 genomes, we calculated the time between the date of index case and the tMRCA after rejection sampling (Fig. 3) and multiplied that time (in years) by the corresponding mean substitution rate from the posterior sample from BEAST.

## Acknowledgements

We thank the patients and healthcare workers who made the collection of this global viral data set possible and all those who made viral genomic data available for analysis. We thank Jennifer Havens and Michael Sanderson for fruitful conversations and insight.

## Author Contributions

Conceptualization: JOW

Methodology: JP,NM,MW,KS,JOW

Software: JP,NM,JOW

Validation: JP,JOW

Formal analysis: JP,JOW

Investigation: JP,JOW

Resources: JOW

Data Curation: JOW

Writing - original draft preparation: JP,JOW

Writing - review and editing: JP,MW,KS,JOW,NM

Visualization: JP,JOW

Supervision: JOW

Project administration: JOW

Funding acquisition: JOW, MW

## Data and materials availability

The BEAST XML files (sequences removed; sequence accession numbers intact), FAVITES JSON files, and GISAID accession numbers for all sequences in this analysis are available at https://github.com/pekarj/SC2_Index_Case.

## Supplementary Materials

Supplementary Results

Tables S1–S4

Figs S1–S10

## Supplementary Materials

### Supplementary Text

#### Relaxed molecular clock

Strict molecular clocks can produce overly-precise tMRCA estimates when evolution occurred under a relaxed clock. Relaxing the molecular clock for SARS-CoV-2 in China using an uncorrelated lognormal distribution of rates (ULD) did produce a slightly wider tMRCA estimate, with a mean of December 6th (95% HPD: November 9th–December 22nd) (Fig. S1) with a comparable rate of 8.45×10^-4^ (95% HPD: 7.05×10^-4^–9.89×10^-4^). However, the standard deviation of the ULD was 0.0009 (95% HPD: 0.00005–0.00013), suggesting strong (strict) clock-like evolution across the SARS-CoV-2 phylogeny in China. Furthermore, 91.7% of the posterior tMRCA estimate post-dated the earliest reported case on 17 November 2019.

#### Phylogenetic rooting

The position of the root in the Bayesian phylodynamic inference under a strict molecular clock was ambiguous. In 63.5% of the posterior trees, the root fell within the basal polytomy of 79 identical genomes, exemplified by the Wuhan-Hu-1 reference genome (GenBank Accession MN908947). This position aligns with the assumption of the NextStrain algorithm (https://nextstrain.org/ncov/global). In 21.8% of posterior trees, the root fell on a branch which led to a single virus: the earliest sampled genome IPBCAMS-WH-01 in 15.0% of trees and another early genome WH01 in 2.4% of trees. This hypothetical root orientation corresponds with the plurality of estimates from Pipes et al. (*20*), though their approach did not make use of the molecular clock. Notably, we find very little support for a root position on branches corresponding to the T28114C (3.9%) or C8782T (1.4%) mutations, as previously suggested by Zhang et al. (*6*).

Rooting configurations were similar when using a relaxed ULD clock. The root fell among viruses with the Wuhan-Hu-1 reference genome sequence in 61.7% of the posterior sample, on the branch leading to PBCAMS-WH-01 in 18.1% of samples, and on the branch leading to WH01 in 3.7% of samples. Again, there was little support for a root position on branches corresponding to the T28114C (2.5%) or C8782T (0.4%) mutations.

#### Time to stable coalescence in simulations

Across the simulated epidemics, a stable coalescence (i.e., non-shifting tMRCA from that point in time until the end of the simulated epidemic) was established after a median of 16 days after the index case was first infected; 95% of simulations reached a stable coalescence by day 53 post-index case infection (Fig. S10A), indicating that the 100-day simulations were sufficient to determine time to stable coalescence. Importantly, the time at which stable coalescence was achieved represents when the tMRCA of the currently circulating lineages is within one day of the tMRCA of the lineages remaining at the end of the simulation. This point in time is when basal lineages ceased to be lost in the coalescent tree, not the tMRCA of the remaining viral lineages (Fig. S10B). At the time a stable coalescence was reached, a median of 16 people had been infected; 95% of simulations reached this stable coalescence after 1194 people had become infected. Recall that the empirical phylogenetic analysis from the Chinese epidemic appears to have reached a stable coalescence by 01 January 2020, when around 1000 people are believed to have been infected (*22*).

#### Number of hospitalized individuals through time

There has also been speculation that hospitals in Hubei province were inundated with COVID-19 patients in October and November 2019 (*36*). However, our primary simulation analysis conditioning on the first case being identified by 17 November 2019 suggests that an increase in hospitalizations due to COVID-19 would not have been notable until mid-to late-December. We note that our estimate of 0 to 36 hospitalizations as of 01 January 2020 in the primary analysis is less than the 42 hospitalizations that had been previously reported (*7*). However, this value is contained within the 95% HPD of many robustness analyses (Fig. S8). Further, it is important to acknowledge that our model is based on parameters from Hao et al. during the period between 01 January and 22 January 2020, which occurred after SARS-CoV-2 was discovered and while there was still limited understanding of COVID-19 (*22*). Therefore, we have little confidence in the precision of our estimates regarding the number of hospitalized patients in late-2019. Nonetheless, even across the various robustness analyses we explored, we never observed a substantial number of hospitalized patients in November 2019, even in the 99% extreme of our estimates (Fig. S8).

**Table S1.** Simulation parameters, with all parameters except for *b* based on Hao et al. (*22*). The *b* value listed is for the primary analysis.

| Parameter | Meaning | Value |
| --- | --- | --- |
| $b$ | Transmission rate of ascertained cases | 0.385 |
| $r$ | Ascertainment rate | 0.15 |
| $a$ | Ratio of transmission of unascertained to ascertained cases | 0.55 |
| $D_e$ | Latent period (days) | 2.9 |
| $D_p$ | Presymptomatic infectious period (days) | 2.3 |
| $D_i$ | Symptomatic infectious period (days) | 2.9 |
| $D_q$ | Duration from illness onset to isolation/hospitalization (days) | 21 |
| $D_h$ | Isolation/hospitalization period (days) | 30 |

**Table S2.** Parameterization and success/failure of compartmental epidemic simulations in primary analysis and robustness analyses.

| Parameter | Primary analysis | Robustness analyses |  |  |  |
| --- | --- | --- | --- | --- | --- |
|  |  | Less infectious | More infectious | Fewer connections | More connections |
| Average number of edges per node (degree) | 16 | 16 | 16 | 10 | 26 |
| Infectiousness (*b*) | 0.385 | 0.300 | 0.550 | 0.385 | 0.385 |
| Doubling time of established simulations<sup>a</sup> | 4.1<br>(2.7-6.7) | 5.3<br>(3.6-7.5) | 3.1<br>(1.6-5.1) | 4.2<br>(2.7-7.0) | 4.2<br>(2.7-7.1) |
| Percentage failed | 70.3 | 80.5 | 53.6 | 69.4 | 68.3 |
| Days to extinction of failed simulations<sup>a</sup> | 7.9<br>(1.5-40.4) | 8.3<br>(1.6-45.7) | 7.2<br>(1.4-34.2) | 8.0<br>(1.5-42.7) | 7.6<br>(1.4-40.1) |
| Number of infections of failed simulations<sup>a</sup> | 1<br>(1-9) | 1<br>(1-11) | 1<br>(1-7) | 1<br>(1-9) | 1<br>(1-9) |
| Maximum number of infections in failed simulations | 44 | 69 | 20 | 29 | 33 |

<sup>a</sup>Median and 95% range in parentheses

**Table S3.**
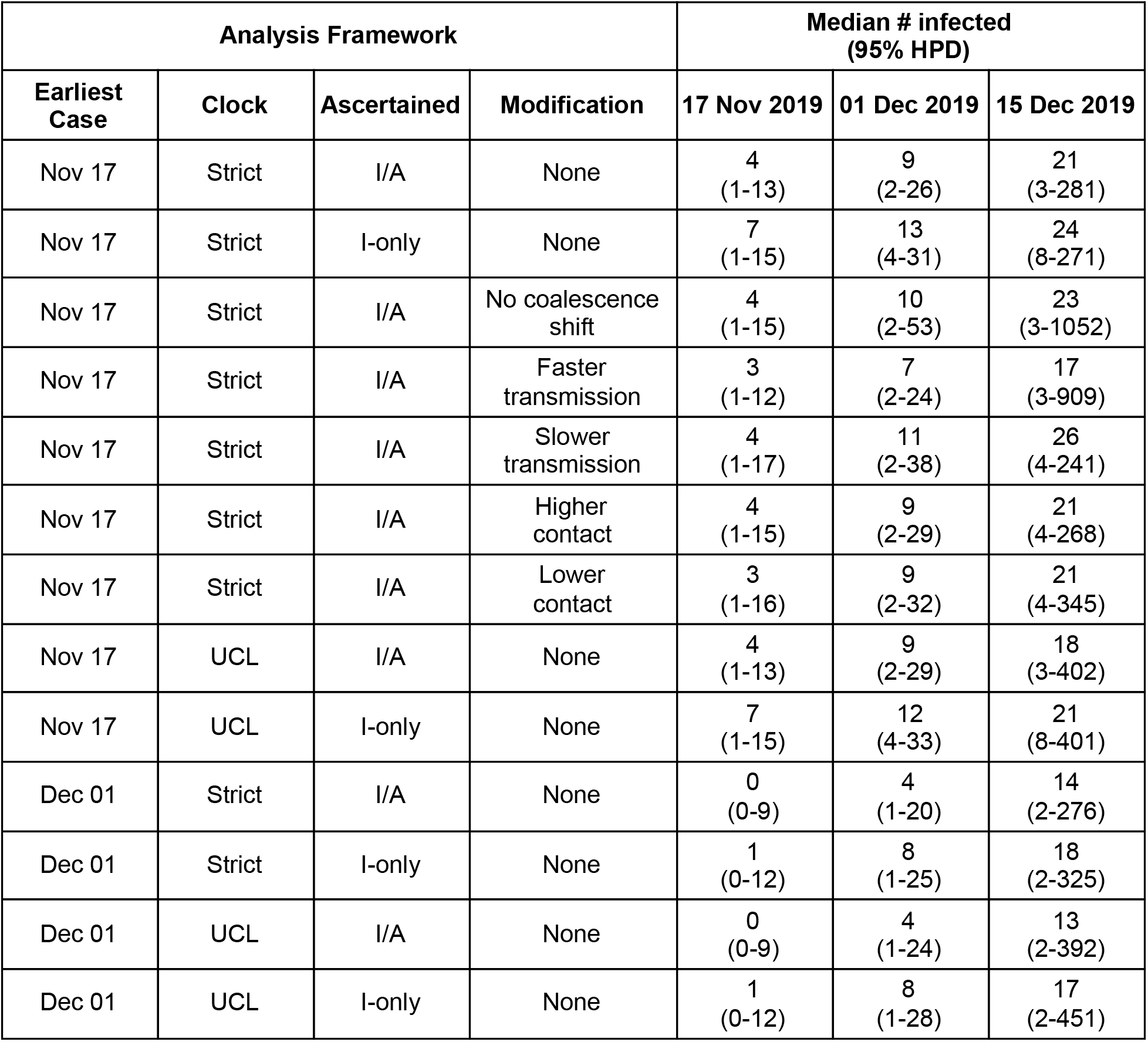
Number infected estimates for primary (first row) and robustness analyses, with the number infected reported on 17 November, 01 December, and 15 December 2019.

**Table S4.** FAVITES parameter values for the primary analysis. Rate values and end time are adjusted for years and density of contact network as needed.

| Parameter<sup>a</sup> | Interpretation | Value |
| --- | --- | --- |
| *num_cn_nodes* | Total number of nodes in network | 100,000 |
| *num_seeds* | Number of seeds (individuals in compartments E,P,I,A) at time 0 | 1 |
| *saphire_freq_s* | Frequency of S individuals at time 0 | 99,999 |
| *saphire_freq_e* | Frequency of E individuals at time 0 | 1 |
| *saphire_freq_p* | Frequency of P individuals at time 0 | 0 |
| *saphire_freq_i* | Frequency of I individuals at time 0 | 0 |
| *saphire_freq_a* | Frequency of A individuals at time 0 | 0 |
| *saphire_freq_h* | Frequency of H individuals at time 0 | 0 |
| *saphire_freq_r* | Frequency of R individuals at time 0 | 0 |
| *saphire_s_to_e_seed* | Target infection rate (S→E) from outside the contact network (i.e., seed infection) | 0 |
| *saphire_e_to_p* | Target rate of becoming presymptomatic (E→P) | 365/2.9 |
| *saphire_p_to_i* | Target rate of becoming ascertained (P→I) | 0.15*365/2.3 |
| *saphire_p_to_a* | Target rate of becoming unascertained (P→A) | (1-0.15)*365/2.3 |
| *saphire_i_to_h* | Target rate of becoming hospitalized (I→H) | 365/21 |
| *saphire_i_to_r* | Target rate of becoming removed/recovered (I→R) | 365/2.9 |
| *saphire_a_to_r* | Target rate of becoming removed/recovered (A→R) | 365/2.9 |
| *saphire_h_to_r* | Target rate of becoming removed/recovered (H→R) | 365/30 |
| *saphire_s_to_e_by_p* | Target infection rate (S→E) by P | 0.55*0.385*365/16 |
| *saphire_s_to_e_by_i* | Target infection rate (S→E) by I | 0.385*365/16 |
| *saphire_s_to_e_by_a* | Target infection rate (S→E) by A | 0.55*0.385*365/16 |
| *end_time* | Time at which to end the transmission simulation | 100/365 |
| *vts_model* | Intrahost viral population growth model to use | constant |
| *vts_growthRate* | Target effective population size growth rate (used in exponential and logistic models) | 0 |
| *vts_max_attempts* | Maximum number of attempts to coalesce a single tree (e.g. 100) before FAVITES kills VirusTreeSimulator | 1000 |
| *vts_n0* | Target effective population size at time zero (used in all models) | 1 |
| *vts_t50* | Time point, relative to the time of infection in backwards time, at which the population is equal to half its final asymptotic value, in the logistic model | -99999 |

<sup>a</sup>FAVITES module choices and parameter values for robustness analyses can be found in the json files at https://github.com/pekarj/SC2_Index_Case.

**Fig. S1.**
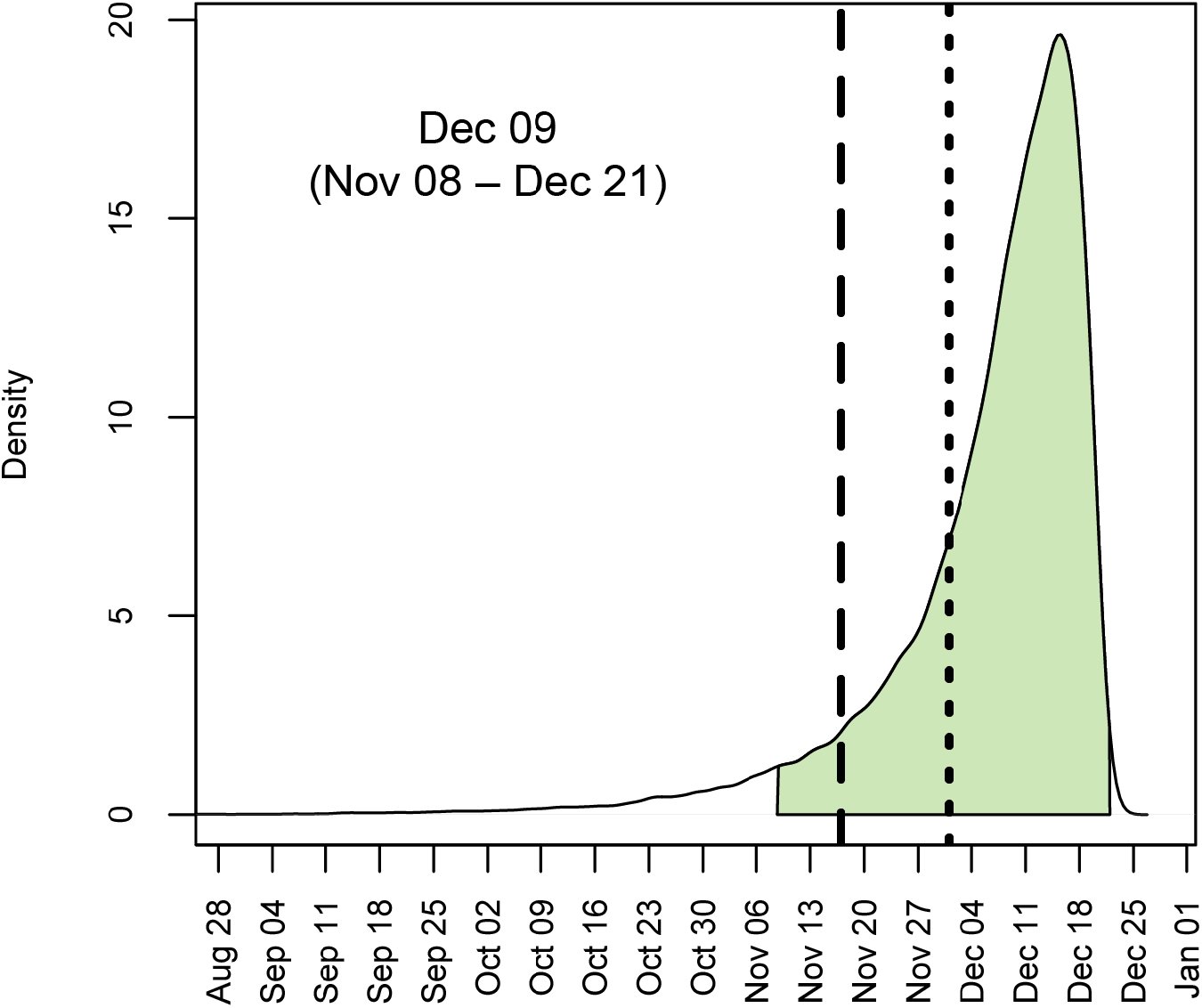
Posterior distribution for the tMRCA of 583 sampled SARS-CoV-2 genomes circulating in China between December 2019 and April 2020 using a relaxed molecular clock (ULD). Shaded area denotes 95% HPD. Long-dashed line is 17 November 2019, and short-dashed line is 01 December 2019.

**Fig. S2.**
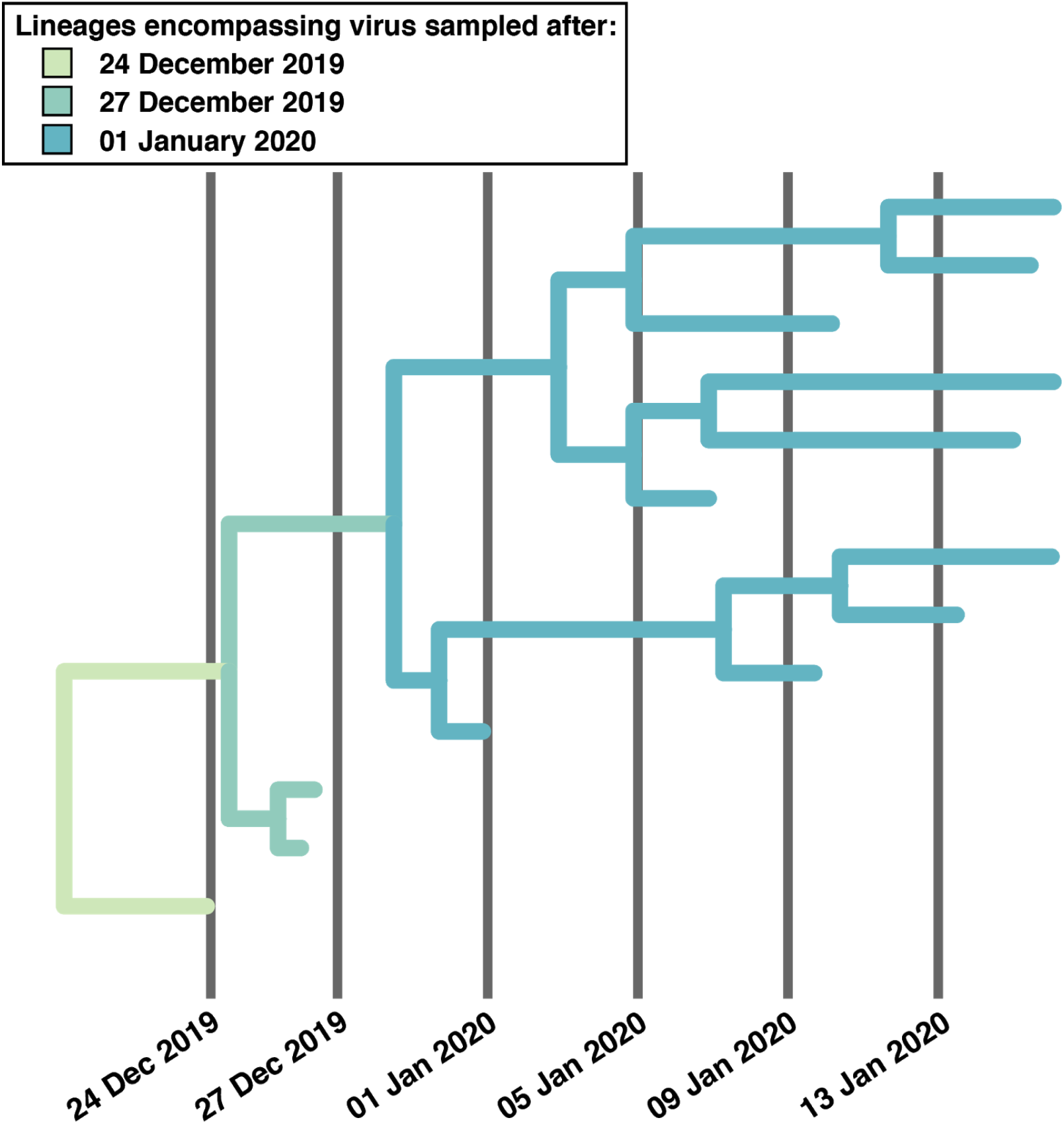
Example of how coalescence of all sampled genomes can shift forward when analyzing viral genomes sampled in late-December 2019 and January 2020 as basal lineages are lost and cease to propagate. In this example, viruses sampled after 01 January 2020 have a stable tMRCA.

**Fig. S3.**
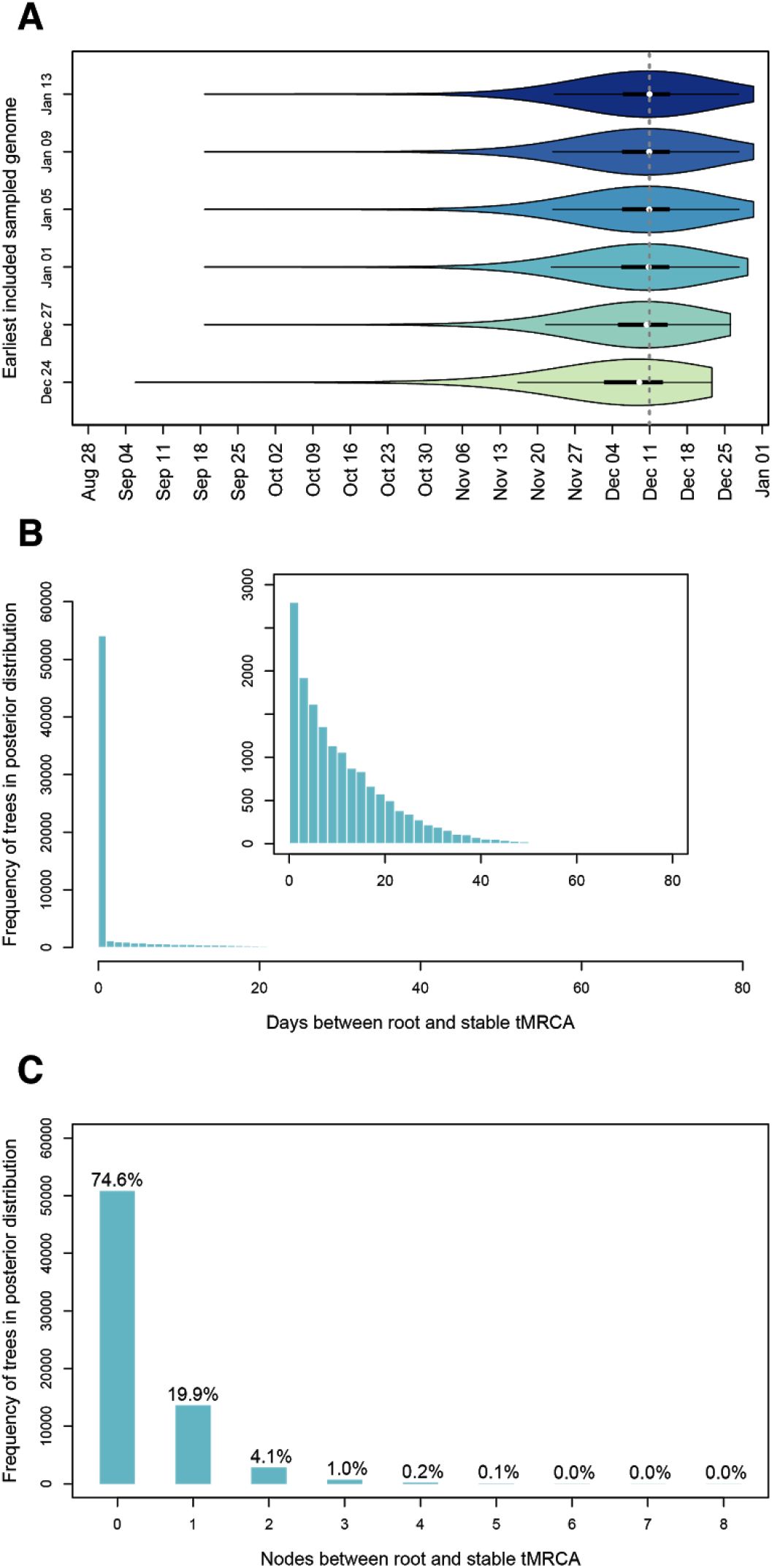
Shifting of tMRCA in empirical molecular clock analyses. (A) Violin plots of tMRCA estimates upon excluding the earliest sampled genomes between 24 December 2019 and 13 January 2020 (darker colors exclude progressively more early sampled genomes). Mean tMRCAs are depicted with white dots. The dashed grey line represents the mean estimate when including only genotypes sampled on 13 January 2020 or later. (B) The number of days between the tMRCA of all genomes (i.e., root) and the tMRCA of genomes sampled on or after 01 January 2020. Inset excludes BEAST samples with <1 day shift. (C) Number of nodes between the tMRCA of all genomes (i.e., root) and the tMRCA of genomes sampled on or after 01 January 2020. Percentage denotes the number of BEAST samples represented in column.

**Fig. S4.**
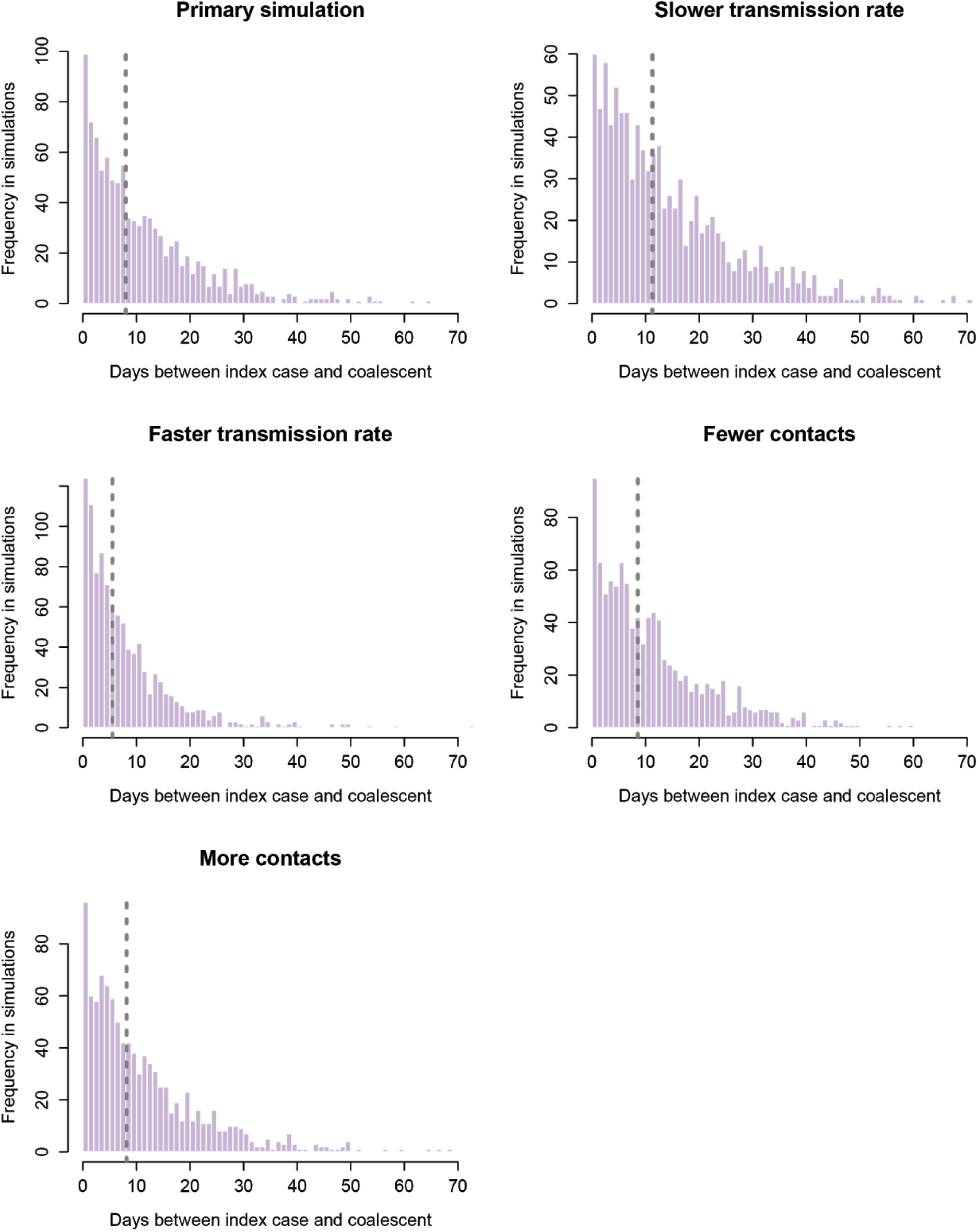
FAVITES robustness analyses. Number of days between the index case the coalescence when simulated transmission rate is slower and faster and when the contact network has fewer (median=10) and more (median=26) contacts.

**Fig. S5.**
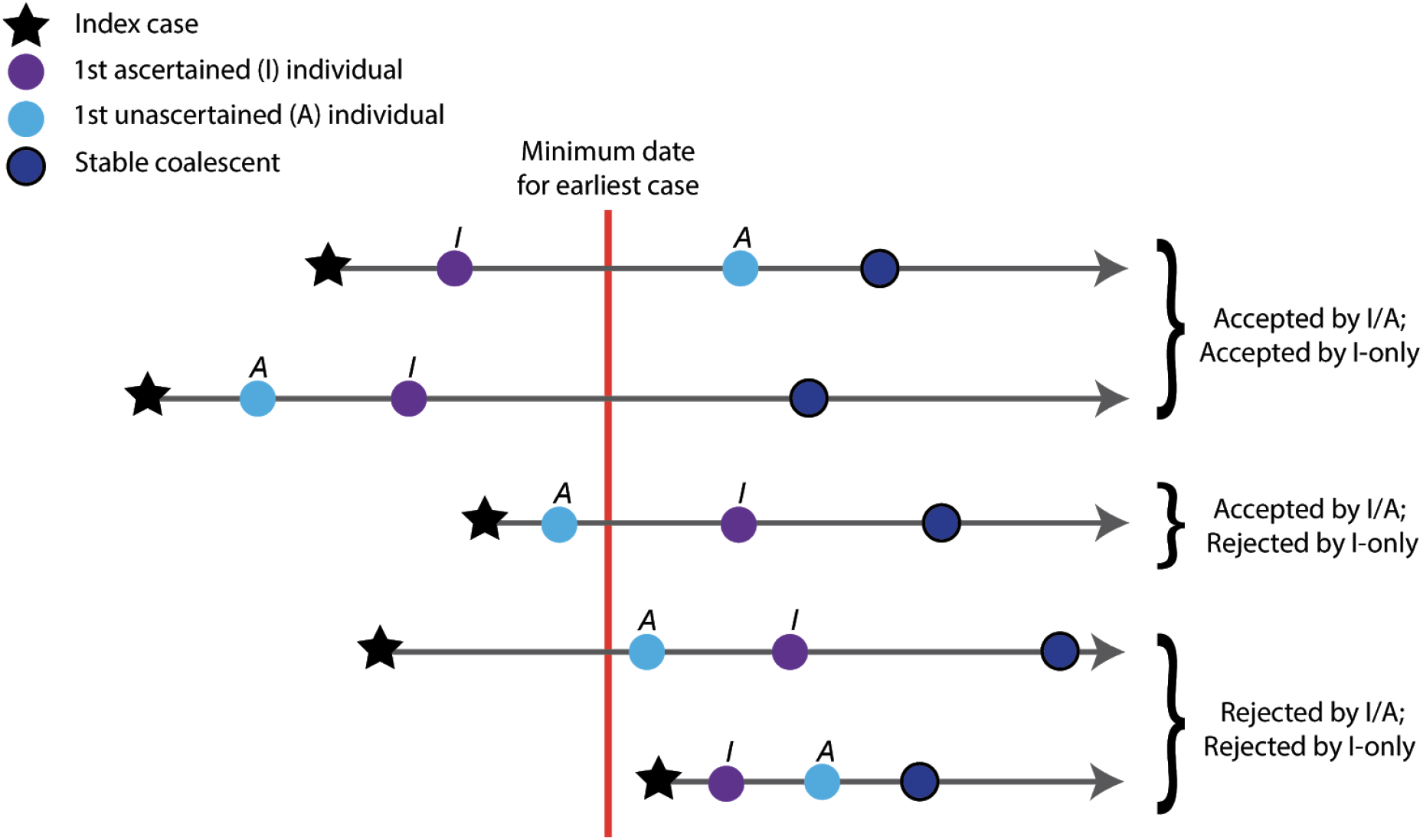
Illustration of the interplay rejection sampling scheme and the SAPHIRE compartmental model. The top two examples would be accepted by both I/A and I-only rejection sampling, because the first ascertained (I) case came before the minimum date for the earliest case In contrast, the middle example would only be accepted by I/A sampling, because the first unascertained (A) case came before the minimum date for the earliest case, but the first ascertained (I) case did not. The bottom two examples would be always be rejected, because both the first ascertained and unascertained cases came after the minimum date for the earliest case Note that coalescence can happen any time after the index case is first infected, and there is no requirement for coalescence to occur after the first ascertained and unascertained individuals. The index case begins in the exposed (E) compartment and can be the individual that first transitions into the A or I compartments; though other subsequently infected individuals can progress to the A or I compartments before index case.

**Fig. S6.**
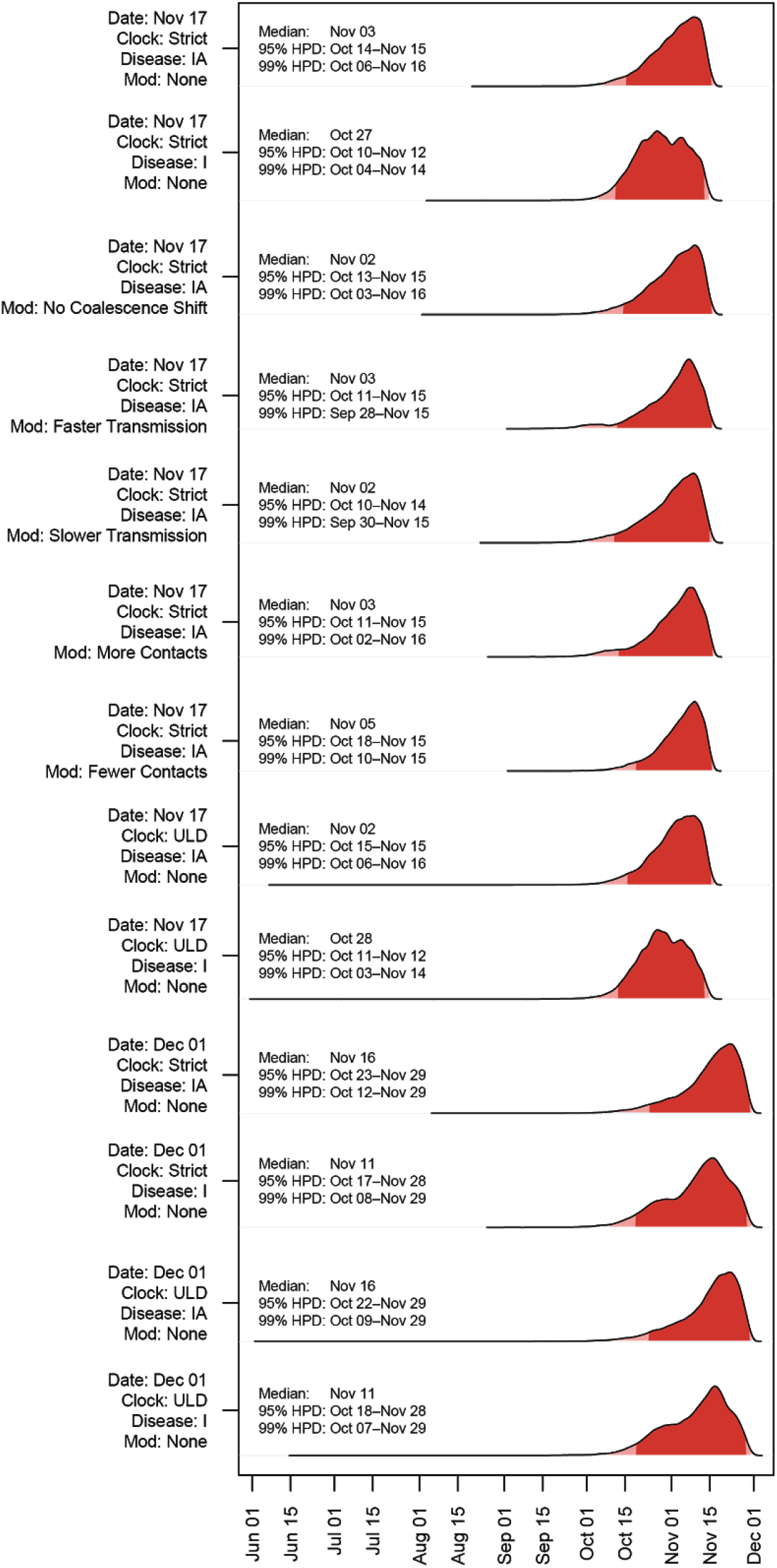
Robustness analysis for timing of SARS-CoV-2 index case in Hubei province. The 95% HPD is shown in dark red, and the 99% HPD is shown in light red. Primary analysis is shown on top. Dates indicates the minimum bound for rejection sampling (Nov 17 or Dec 01). Clock indicates whether the tMRCA was inferred using a strict or relaxed (ULD) clock. Disease denotes which stage of infection in the SAPHIRE model (IA, Ascertained or Unascertained; or I, Ascertained only) must have been reached by the given date for rejection sampling. ‘Mod’ denotes if any robustness modifications were explored.

**Fig. S7.**
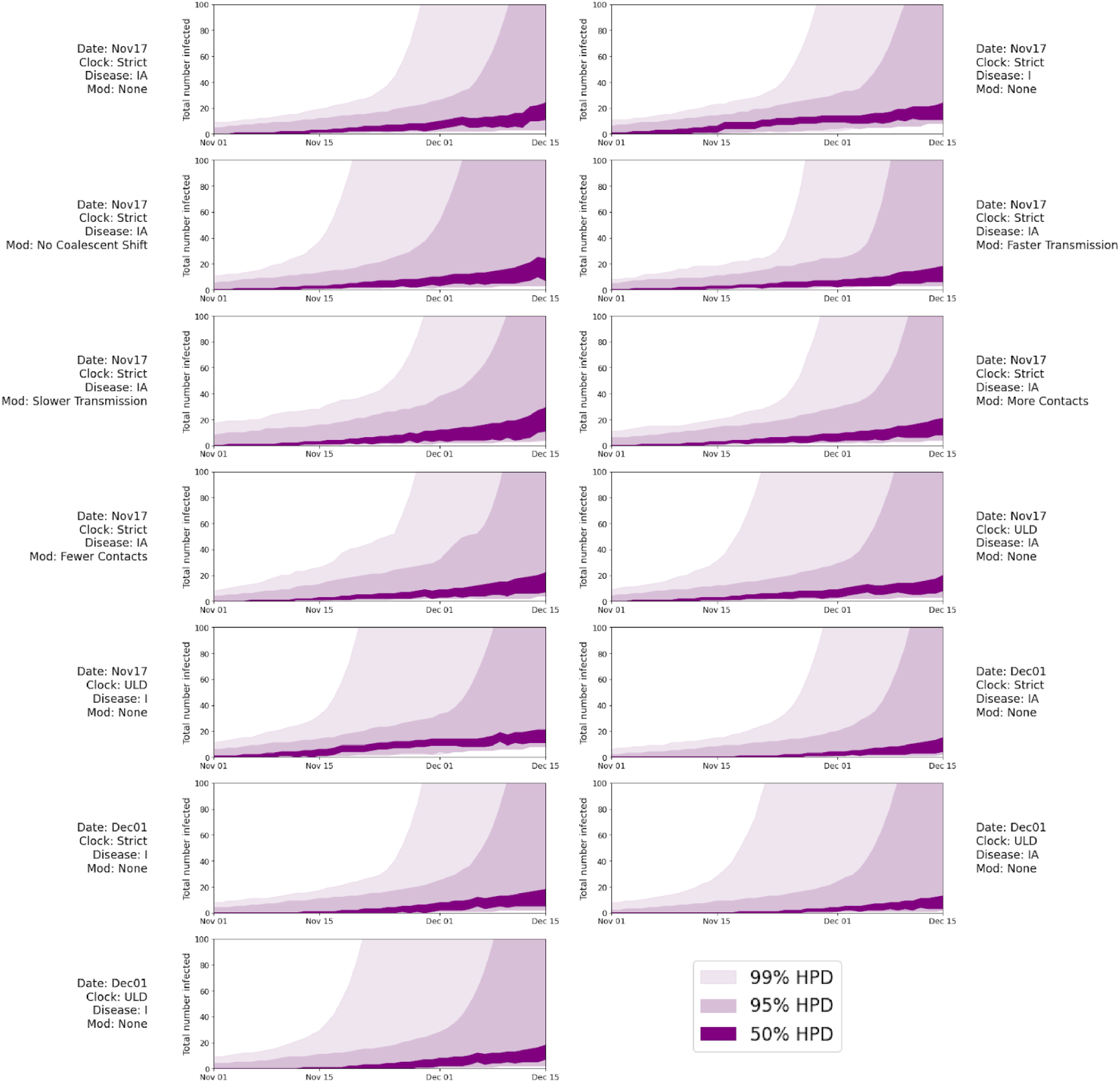
Robustness analysis for the number of people infected SARS-CoV-2 based on the SAPHIRE model in late 2019. Innermost shading is 50% HPD, middle shading is 95% HPD, and outer shading is 99% HPD. Dates indicates the minimum bound for rejection sampling (Nov 17 or Dec 01). Clock indicates whether the tMRCA was inferred using a strict or relaxed (ULD) clock. Disease denotes which stage of infection in the SAPHIRE model (IA, Ascertained or Unascertained; or I, Ascertained only) must have been reached by the given date for rejection sampling.

**Fig. S8.**
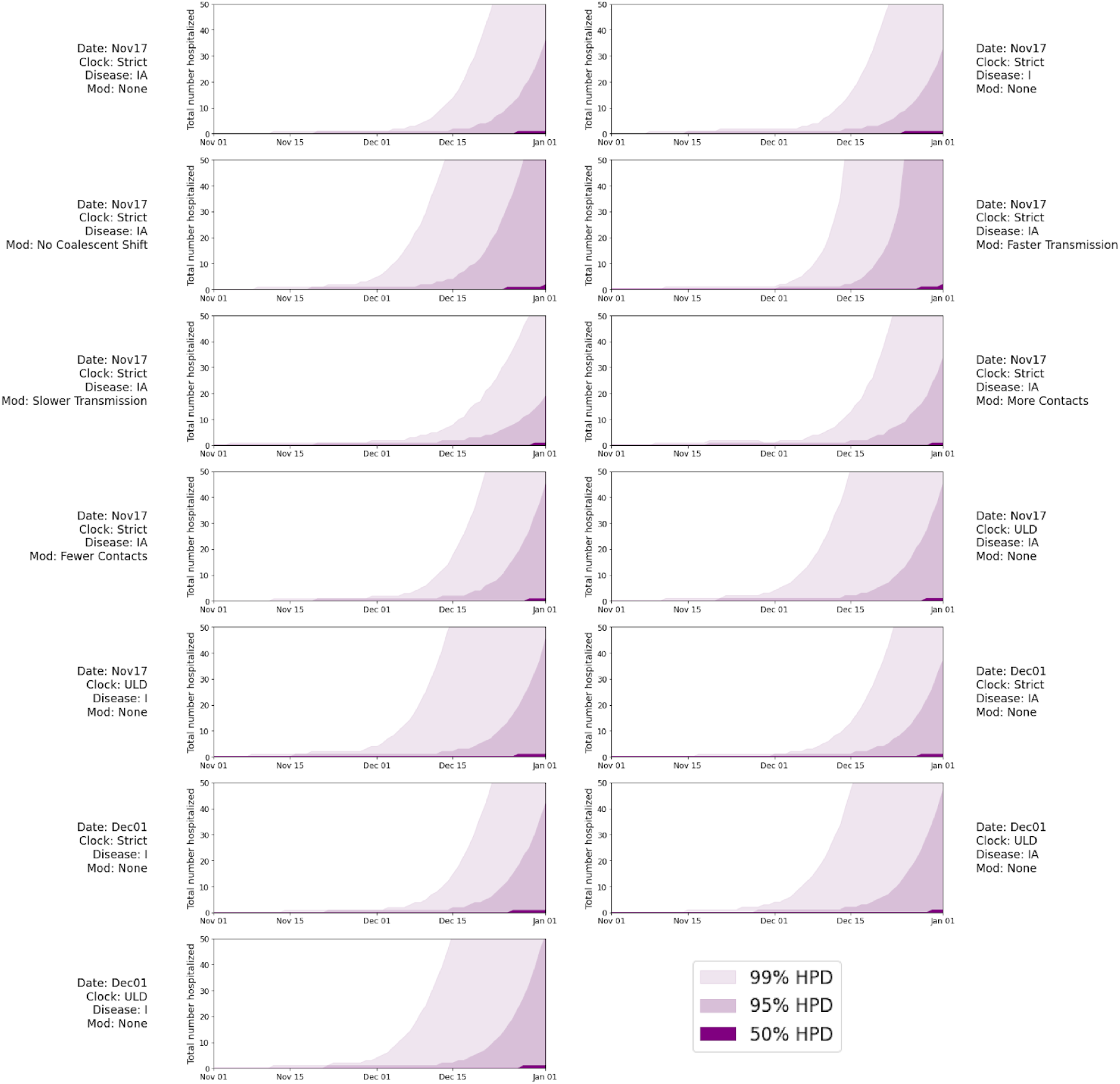
Number of people hospitalized with SARS-CoV-2 based on the SAPHIRE model in late 2019. Innermost shading is 50% HPD, middle shading is 95% HPD, and outer shading is 99% HPD. Dates indicates the minimum bound for rejection sampling (Nov 17 or Dec 01). Clock indicates whether the tMRCA was inferred using a strict or relaxed (ULD) clock. Disease denotes which stage of infection in the SAPHIRE model (IA, Ascertained or Unascertained; or I, Ascertained only) must have been reached by the given date for rejection sampling.

**Fig. S9.**
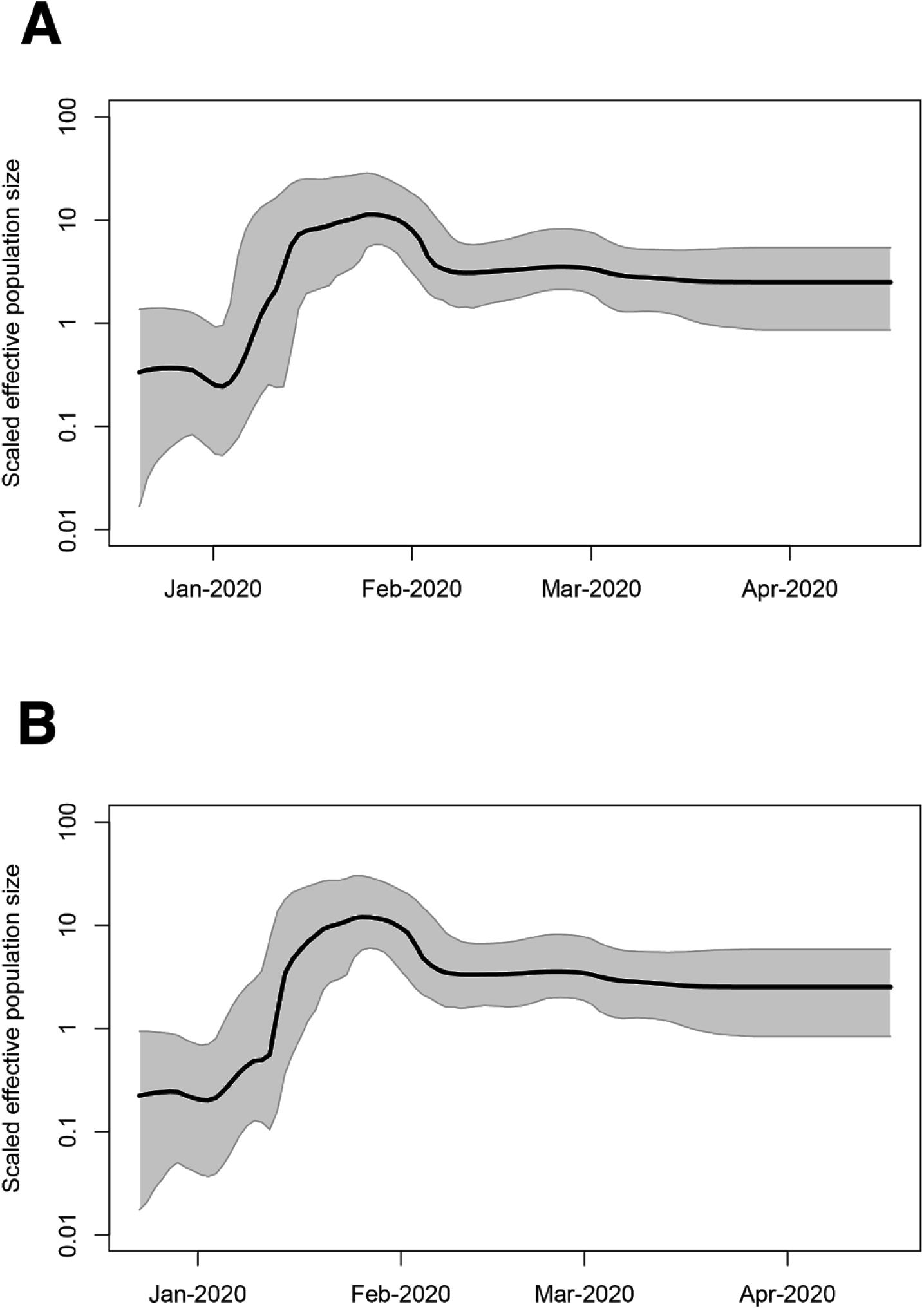
Bayesian skyline plot reconstruction. (A) Strict molecular clock analysis and (B) ULD relaxed molecular clock. Solid line is the median estimate. Shaded area is the 95% HPD.

**Fig. S10.**
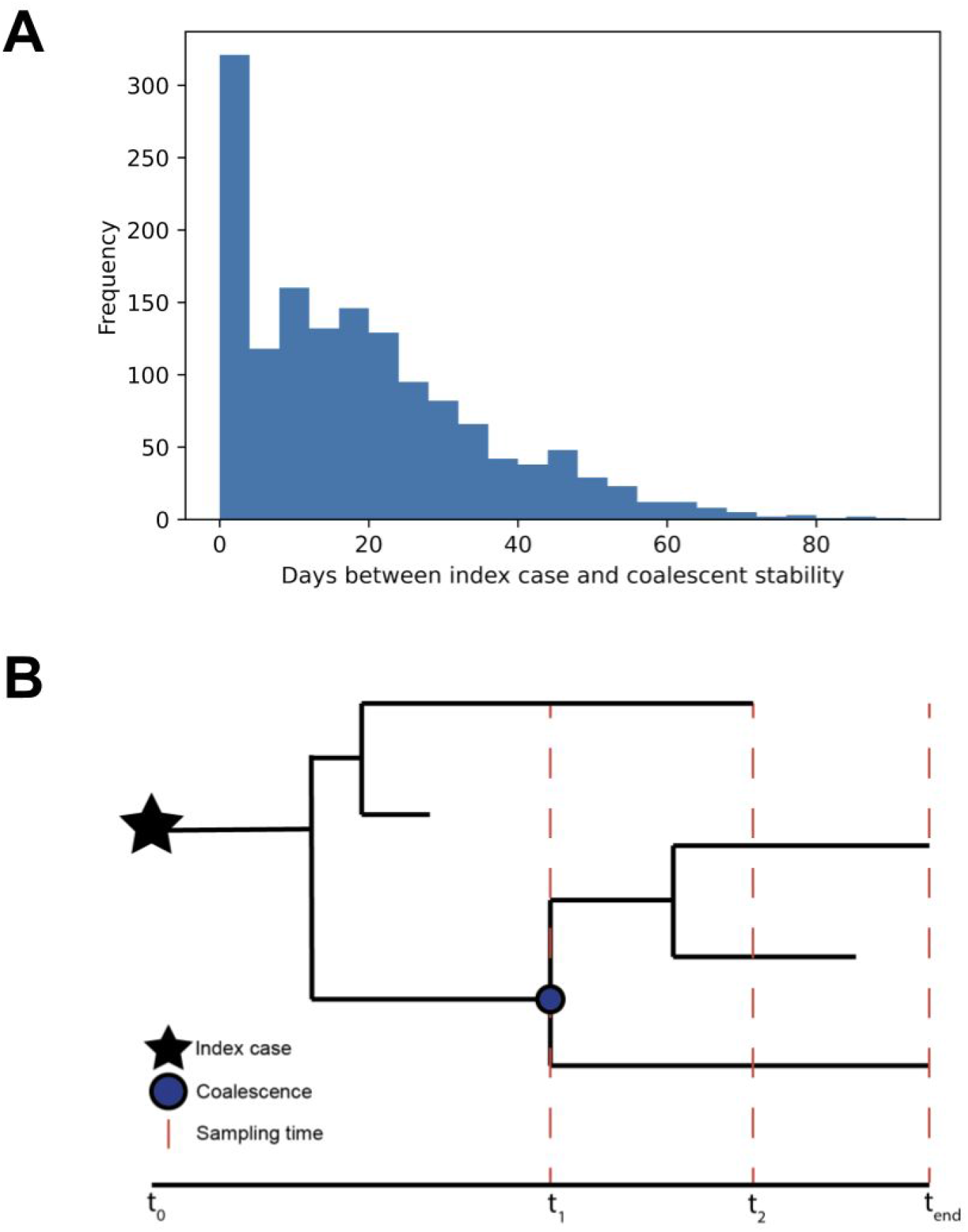
Stable coalesce in forward compartmental simulations. (A) Days between index case infection and time stable coalescence is achieved in primary simulations. (B) Distinction between the date of index case infection (at t_0_), the tMRCA of surviving lineages (at t_1_), and the time at which stable coalescence is achieved when the last basal lineage goes extinct (at t_2_).

## References and Notes

1. N. Zhu, D. Zhang, W. Wang, X. Li, B. Yang, J. Song, X. Zhao, B. Huang, W. Shi, R. Lu, P. Niu, F. Zhan, X. Ma, D. Wang, W. Xu, G. Wu, G. F. Gao, W. Tan, A Novel Coronavirus from Patients with Pneumonia in China, 2019. N. Engl. J. Med. (2020), doi:10.1056/NEJMoa2001017.

2. Q. Li, X. Guan, P. Wu, X. Wang, L. Zhou, Y. Tong, R. Ren, K. S. M. Leung, E. H. Y. Lau, J. Y. Wong, X. Xing, N. Xiang, Y. Wu, C. Li, Q. Chen, D. Li, T. Liu, J. Zhao, M. Liu, W. Tu, C. Chen, L. Jin, R. Yang, Q. Wang, S. Zhou, R. Wang, H. Liu, Y. Luo, Y. Liu, G. Shao, H. Li, Z. Tao, Y. Yang, Z. Deng, B. Liu, Z. Ma, Y. Zhang, G. Shi, T. T. Y. Lam, J. T. Wu, G. F. Gao, B. J. Cowling, B. Yang, G. M. Leung, Z. Feng, Early Transmission Dynamics in Wuhan, China, of Novel Coronavirus-Infected Pneumonia. N. Engl. J. Med. (2020), doi:10.1056/NEJMoa2001316.

3. Report of the WHO-China Joint Mission on Coronavirus Disease 2019 (COVID-19), (available at https://www.who.int/publications-detail-redirect/report-of-the-who-china-joint-mission-on-coronavirus-disease-2019-(covid-19)).

4. H. Fei, X. Yinyin, C. Hui, W. Ni, D. Xin, C. Wei, L. Tao, H. Shitong, S. Miaomiao, C. Mingting, S. Keshavjee, Z. Yanlin, D. P. Chin, L. Jianjun, The impact of the COVID-19 epidemic on tuberculosis control in China. Lancet Reg. Health - West. Pac. 3, 100032 (2020).

5. WHO Director-General’s opening remarks at the media briefing on COVID-19-11 March 2020, (available at https://www.who.int/director-general/speeches/detail/who-director-general-s-opening-remarks-at-the-media-briefing-on-covid-19---11-march-2020).

6. X. Zhang, Y. Tan, Y. Ling, G. Lu, F. Liu, Z. Yi, X. Jia, M. Wu, B. Shi, S. Xu, J. Chen, W. Wang, B. Chen, L. Jiang, S. Yu, J. Lu, J. Wang, M. Xu, Z. Yuan, Q. Zhang, X. Zhang, G. Zhao, S. Wang, S. Chen, H. Lu, Viral and host factors related to the clinical outcome of COVID-19. Nature (2020), doi:10.1038/s41586-020-2355-0.

7. C. Huang, Y. Wang, X. Li, L. Ren, J. Zhao, Y. Hu, L. Zhang, G. Fan, J. Xu, X. Gu, Z. Cheng, T. Yu, J. Xia, Y. Wei, W. Wu, X. Xie, W. Yin, H. Li, M. Liu, Y. Xiao, H. Gao, L. Guo, J. Xie, G. Wang, R. Jiang, Z. Gao, Q. Jin, J. Wang, B. Cao, Clinical features of patients infected with 2019 novel coronavirus in Wuhan, China. The Lancet. 395, 497–506 (2020).

8. F. Wu, S. Zhao, B. Yu, Y.-M. Chen, W. Wang, Z.-G. Song, Y. Hu, Z.-W. Tao, J.-H. Tian, Y.-Y. Pei, M.-L. Yuan, Y.-L. Zhang, F.-H. Dai, Y. Liu, Q.-M. Wang, J.-J. Zheng, L. Xu, E. C. Holmes, Y.-Z. Zhang, A new coronavirus associated with human respiratory disease in China. Nature. 579, 265–269 (2020).

9. R. Lu, X. Zhao, J. Li, P. Niu, B. Yang, H. Wu, W. Wang, H. Song, B. Huang, N. Zhu, Y. Bi, X. Ma, F. Zhan, L. Wang, T. Hu, H. Zhou, Z. Hu, W. Zhou, L. Zhao, J. Chen, Y. Meng, J. Wang, Y. Lin, J. Yuan, Z. Xie, J. Ma, W. J. Liu, D. Wang, W. Xu, E. C. Holmes, G. F. Gao, G. Wu, W. Chen, W. Shi, W. Tan, Genomic characterisation and epidemiology of 2019 novel coronavirus: implications for virus origins and receptor binding. Lancet Lond. Engl. 395, 565–574 (2020).

10. Coronavirus: China’s first confirmed Covid-19 case traced back to November 17 | South China Morning Post, (available at https://www.scmp.com/news/china/society/article/3074991/coronavirus-chinas-first-confirmed-covid-19-case-traced-back).

11. S. Duchene, L. Featherstone, M. Haritopoulou-Sinanidou, A. Rambaut, P. Lemey, G. Baele, Temporal signal and the phylodynamic threshold of SARS-CoV-2. Virus Evol., doi:10.1093/ve/veaa061.

12. A. Rambaut, Phylodynamic Analysis | 176 genomes | 6 Mar 2020. Virological (2020), (available at https://virological.org/t/phylodynamic-analysis-176-genomes-6-mar-2020/356).

13. K. G. Andersen, A. Rambaut, W. I. Lipkin, E. C. Holmes, R. F. Garry, The proximal origin of SARS-CoV-2. Nat. Med. 26, 450–452 (2020).

14. N. R. Faria, A. Rambaut, M. A. Suchard, G. Baele, T. Bedford, M. J. Ward, A. J. Tatem, J. D. Sousa, N. Arinaminpathy, J. Pépin, D. Posada, M. Peeters, O. G. Pybus, P. Lemey, The early spread and epidemic ignition of HIV-1 in human populations. Science. 346, 56–61 (2014).

15. M. Worobey, M. Gemmel, D. E. Teuwen, T. Haselkorn, K. Kunstman, M. Bunce, J.-J. Muyembe, J.-M. M. Kabongo, R. M. Kalengayi, E. Van Marck, M. T. P. Gilbert, S. M. Wolinsky, Direct evidence of extensive diversity of HIV-1 in Kinshasa by 1960. Nature. 455, 661–664 (2008).

16. B. F. Keele, F. V. Heuverswyn, Y. Li, E. Bailes, J. Takehisa, M. L. Santiago, F. Bibollet-Ruche, Y. Chen, L. V. Wain, F. Liegeois, S. Loul, E. M. Ngole, Y. Bienvenue, E. Delaporte, J. F. Y. Brookfield, P. M. Sharp, G. M. Shaw, M. Peeters, B. H. Hahn, Chimpanzee Reservoirs of Pandemic and Nonpandemic HIV-1. Science. 313, 523–526 (2006).

17. T. Bedford, A. L. Greninger, P. Roychoudhury, L. M. Starita, M. Famulare, M.-L. Huang, A. Nalla, G. Pepper, A. Reinhardt, H. Xie, L. Shrestha, T. N. Nguyen, A. Adler, E. Brandstetter, S. Cho, D. Giroux, P. D. Han, K. Fay, C. D. Frazar, M. Ilcisin, K. Lacombe, J. Lee, A. Kiavand, M. Richardson, T. R. Sibley, M. Truong, C. R. Wolf, D. A. Nickerson, M. J. Rieder, J. A. Englund, T. S. F. S. Investigators, J. Hadfield, E. B. Hodcroft, J. Huddleston, L. H. Moncla, N. F. Müller, R. A. Neher, X. Deng, W. Gu, S. Federman, C. Chiu, J. S. Duchin, R. Gautom, G. Melly, B. Hiatt, P. Dykema, S. Lindquist, K. Queen, Y. Tao, A. Uehara, S. Tong, D. MacCannell, G. L. Armstrong, G. S. Baird, H. Y. Chu, J. Shendure, K. R. Jerome, Cryptic transmission of SARS-CoV-2 in Washington state. Science (2020), doi:10.1126/science.abc0523.

18. Q. Bi, Y. Wu, S. Mei, C. Ye, X. Zou, Z. Zhang, X. Liu, L. Wei, S. A. Truelove, T. Zhang, W. Gao, C. Cheng, X. Tang, X. Wu, Y. Wu, B. Sun, S. Huang, Y. Sun, J. Zhang, T. Ma, J. Lessler, T. Feng, Epidemiology and transmission of COVID-19 in 391 cases and 1286 of their close contacts in Shenzhen, China: a retrospective cohort study. Lancet Infect. Dis. 20, 911–919 (2020).

19. M. A. Suchard, P. Lemey, G. Baele, D. L. Ayres, A. J. Drummond, A. Rambaut, Bayesian phylogenetic and phylodynamic data integration using BEAST 1.10. Virus Evol. 4 (2018), doi:10.1093/ve/vey016.

20. L. Pipes, H. Wang, J. P. Huelsenbeck, R. Nielsen, “Assessing uncertainty in the rooting of the SaRS-CoV-2 phylogeny” (preprint, Evolutionary Biology, 2020), doi:10.1101/2020.06.19.160630.

21. N. Moshiri, M. Ragonnet-Cronin, J. O. Wertheim, S. Mirarab, FAVITES: simultaneous simulation of transmission networks, phylogenetic trees and sequences. Bioinformatics. 35, 1852–1861 (2019).

22. X. Hao, S. Cheng, D. Wu, T. Wu, X. Lin, C. Wang, Reconstruction of the full transmission dynamics of COVID-19 in Wuhan. Nature, 1–7 (2020).

23. S. Kumar, Q. Tao, S. Weaver, M. Sanderford, M. A. Caraballo-Ortiz, S. Sharma, S. L. K. Pond, S. Miura, “An evolutionary portrait of the progenitor SARS-CoV-2 and its dominant offshoots in COVID-19 pandemic” (preprint, Evolutionary Biology, 2020), doi:10.1101/2020.09.24.311845.

24. R. Laxminarayan, B. Wahl, S. R. Dudala, K. Gopal, C. Mohan, S. Neelima, K. S. J. Reddy, J. Radhakrishnan, J. A. Lewnard, Epidemiology and transmission dynamics of COVID-19 in two Indian states. Science (2020), doi:10.1126/science.abd7672.

25. D. C. Adam, P. Wu, J. Y. Wong, E. H. Y. Lau, T. K. Tsang, S. Cauchemez, G. M. Leung, B. J. Cowling, Clustering and superspreading potential of SARS-CoV-2 infections in Hong Kong. Nat. Med., 1–6 (2020).

26. A. Endo, S. Abbott, A. J. Kucharski, S. Funk, Estimating the overdispersion in COVID-19 transmission using outbreak sizes outside China. Wellcome Open Res. 5 (2020), doi:10.12688/wellcomeopenres.15842.3.

27. X.-K. Xu, X. F. Liu, Y. Wu, S. T. Ali, Z. Du, P. Bosetti, E. H. Y. Lau, B. J. Cowling, L. Wang, Reconstruction of Transmission Pairs for Novel Coronavirus Disease 2019 (COVID-19) in Mainland China: Estimation of Superspreading Events, Serial Interval, and Hazard of Infection. Clin. Infect. Dis., doi:10.1093/cid/ciaa790.

28. M. Worobey, J. Pekar, B. B. Larsen, M. I. Nelson, V. Hill, J. B. Joy, A. Rambaut, M. A. Suchard, J. O. Wertheim, P. Lemey, The emergence of SARS-CoV-2 in Europe and North America. Science (2020), doi:10.1126/science.abc8169.

29. L. du Plessis, J. T. McCrone, A. E. Zarebski, V. Hill, C. Ruis, B. Gutierrez, J. Raghwani, J. Ashworth, R. Colquhoun, T. R. Connor, N. R. Faria, B. Jackson, N. J. Loman, A. O’Toole, S. M. Nicholls, K. V. Parag, E. Scher, T. I. Vasylyeva, E. M. Volz, A. Watts, I. I. Bogoch, K. Khan, C.-U. Consortium, D. Aanensen, M. U. G. Kraemer, A. Rambaut, O. Pybus, medRxiv, in press, doi:10.1101/2020.10.23.20218446.

30. A. Deslandes, V. Berti, Y. Tandjaoui-Lambotte, C. Alloui, E. Carbonnelle, J. R. Zahar, S. Brichler, Y. Cohen, SARS-CoV-2 was already spreading in France in late December 2019. Int. J. Antimicrob. Agents. 55, 106006 (2020).

31. X. Deng, W. Gu, S. Federman, L. du Plessis, O. G. Pybus, N. R. Faria, C. Wang, G. Yu, B. Bushnell, C.-Y. Pan, H. Guevara, A. Sotomayor-Gonzalez, K. Zorn, A. Gopez, V. Servellita, E. Hsu, S. Miller, T. Bedford, A. L. Greninger, P. Roychoudhury, L. M. Starita, M. Famulare, H. Y. Chu, J. Shendure, K. R. Jerome, C. Anderson, K. Gangavarapu, M. Zeller, E. Spencer, K. G. Andersen, D. MacCannell, C. R. Paden, Y. Li, J. Zhang, S. Tong, G. Armstrong, S. Morrow, M. Willis, B. T. Matyas, S. Mase, O. Kasirye, M. Park, G. Masinde, C. Chan, A. T. Yu, S. J. Chai, E. Villarino, B. Bonin, D. A. Wadford, C. Y. Chiu, Genomic surveillance reveals multiple introductions of SARS-CoV-2 into Northern California. Science. 369, 582–587 (2020).

32. CDCMMWR, Evidence for Limited Early Spread of COVID-19 Within the United States, January-February 2020. MMWR Morb. Mortal. Wkly. Rep. 69 (2020), doi:10.15585/mmwr.mm6922e1.

33. G. Chavarria-Miró, E. Anfruns-Estrada, S. Guix, M. Paraira, B. Galofré, G. Sáanchez, R. Pintó, A. Bosch, “Sentinel surveillance of SARS-CoV-2 in wastewater anticipates the occurrence of COVID-19 cases” (preprint, Epidemiology, 2020), doi:10.1101/2020.06.13.20129627.

34. G. Fongaro, P. H. Stoco, D. S. M. Souza, E. C. Grisard, M. E. Magri, P. Rogovski, M. A. Schorner, F. H. Barazzetti, A. P. Christoff, L. F. V. de Oliveira, M. L. Bazzo, G. Wagner, M. Hernandez, D. Rodriguez-Lazaro, medRxiv, in press, doi:10.1101/2020.06.26.20140731.

35. G. L. Rosa, P. Mancini, G. B. Ferraro, C. Veneri, M. Iaconelli, L. Bonadonna, L. Lucentini, E. Suffredini, medRxiv, in press, doi:10.1101/2020.06.25.20140061.

36. E. O. Nsoesie, B. Rader, Y. L. Barnoon, L. Goodwin, J. Brownstein, Analysis of hospital traffic and search engine data in Wuhan China indicates early disease activity in the Fall of 2019 (2020) (available at https://dash.harvard.edu/handle/1/42669767).

37. B. Q. Minh, H. A. Schmidt, O. Chernomor, D. Schrempf, M. D. Woodhams, A. von Haeseler, R. Lanfear, IQ-TREE 2: New Models and Efficient Methods for Phylogenetic Inference in the Genomic Era. Mol. Biol. Evol. 37, 1530–1534 (2020).

38. A. Rambaut, T. T. Lam, L. Max Carvalho, O. G. Pybus, Exploring the temporal structure of heterochronous sequences using TempEst (formerly Path-O-Gen). Virus Evol. 2 (2016), doi:10.1093/ve/vew007.

39. A. Rambaut, A. J. Drummond, D. Xie, G. Baele, M. A. Suchard, Posterior Summarization in Bayesian Phylogenetics Using Tracer 1.7. Syst. Biol. 67, 901–904 (2018).

40. A.-L. Barabási, R. Albert, Emergence of Scaling in Random Networks. Science. 286, 509–512 (1999).

41. J. Mossong, N. Hens, M. Jit, P. Beutels, K. Auranen, R. Mikolajczyk, M. Massari, S. Salmaso, G. S. Tomba, J. Wallinga, J. Heijne, M. Sadkowska-Todys, M. Rosinska, W. J. Edmunds, Social Contacts and Mixing Patterns Relevant to the Spread of Infectious Diseases. PLOS Med. 5, e74 (2008).

42. F. D. Sahneh, A. Vajdi, H. Shakeri, F. Fan, C. Scoglio, GEMFsim: A stochastic simulator for the generalized epidemic modeling framework. J. Comput. Sci. 22, 36–44 (2017).

43. N. Moshiri, TreeSwift: A massively scalable Python tree package. SoftwareX. 11, 100436 (2020).

